# Arp2/3 complex-mediated actin assembly drives microtubule-independent motility and phagocytosis in the evolutionarily divergent amoeba *Naegleria*

**DOI:** 10.1101/2020.05.12.091538

**Authors:** Katrina B. Velle, Lillian K. Fritz-Laylin

**Affiliations:** The University of Massachusetts Amherst, Department of Biology, Amherst, MA 01003

## Abstract

Much of our current understanding of actin-driven phenotypes in eukaryotes has come from the “yeast to human” opisthokont lineage, as well as the related amoebozoa. Outside of these groups lies the genus *Naegleria*, which shared a common ancestor with humans over a billion years ago, and includes the deadly “brain-eating amoeba.” Unlike nearly every other known eukaryotic cell type, *Naegleria* amoebae are thought to lack cytoplasmic microtubules. The absence of microtubules suggests that these amoebae rapidly crawl and phagocytose bacteria using actin alone. Although this makes *Naegleria* a powerful system to probe actin-driven functions in the absence of microtubules, surprisingly little is known about *Naegleria*’s actin cytoskeleton. Here, we use microscopy and genomic analysis to show that *Naegleria* amoebae have an extensive actin cytoskeletal repertoire, complete with nucleators and nucleation promoting factors. *Naegleria* use this cytoskeletal machinery to generate Arp2/3-dependent lamellar protrusions, which correlate with the capacity to migrate and phagocytose bacteria. Because human cells also use Arp2/3-dependent lamellar protrusions for motility and phagocytosis, this work supports an evolutionarily ancient origin for these actin-driven processes and establishes *Naegleria* as a natural model system for studying microtubule-independent cytoskeletal phenotypes.

## INTRODUCTION

Actin is among the most highly expressed proteins in eukaryotic cells, and is thought to have evolved before the origin of eukaryotes ((Akil and Robinson, 2018; Ettema et al., 2011; Spang et al., 2015); and reviewed in (Velle and Fritz-Laylin, 2019)). Actin polymerization drives essential functions that include cell motility, division, phagocytosis, and vesicular trafficking. To accomplish these functions, cells must assemble actin networks precisely when and where they are needed—a complex task that involves layers of regulation (Velle and Fritz-Laylin, 2019). Actin networks are initiated by two main classes of actin nucleators: the Arp/2 complex typically assembles branched actin networks (Mullins et al., 1998), while formins nucleate and elongate linear networks (Breitsprecher and Goode, 2013; Sagot et al., 2002). Actin, the Arp2/3 complex, formins, and many of their regulators are broadly conserved across eukaryotes (Fritz-Laylin et al., 2017a; Kollmar et al., 2012; Pollard and Goldman, 2018; Pruyne, 2017; Velle and Fritz-Laylin, 2019; Veltman and Insall, 2010), but our understanding of cytoskeletal dynamics has come from only a handful of systems, most of which fall within the opisthokonts—the eukaryotic lineage that includes animals and fungi. While studies within this “yeast to human” range have made monumental contributions to cell biology, they do not reflect the diversity of actin cytoskeletal systems within eukarya. Studying additional, diverse organisms has revealed unique cell biology (Craig et al., 2019; Paredez et al., 2014), and provides opportunities to address questions that are intractable using traditionally studied organisms (Russell et al., 2017). In this study, we use the amoeba *Naegleria*—an emerging model system from the heteroloboseans, one of the most evolutionarily distant groups from the opisthokonts (**Fig 1**)—to explore how a cell undergoes crawling motility and phagocytosis in the absence of microtubules.

**Fig 1.**
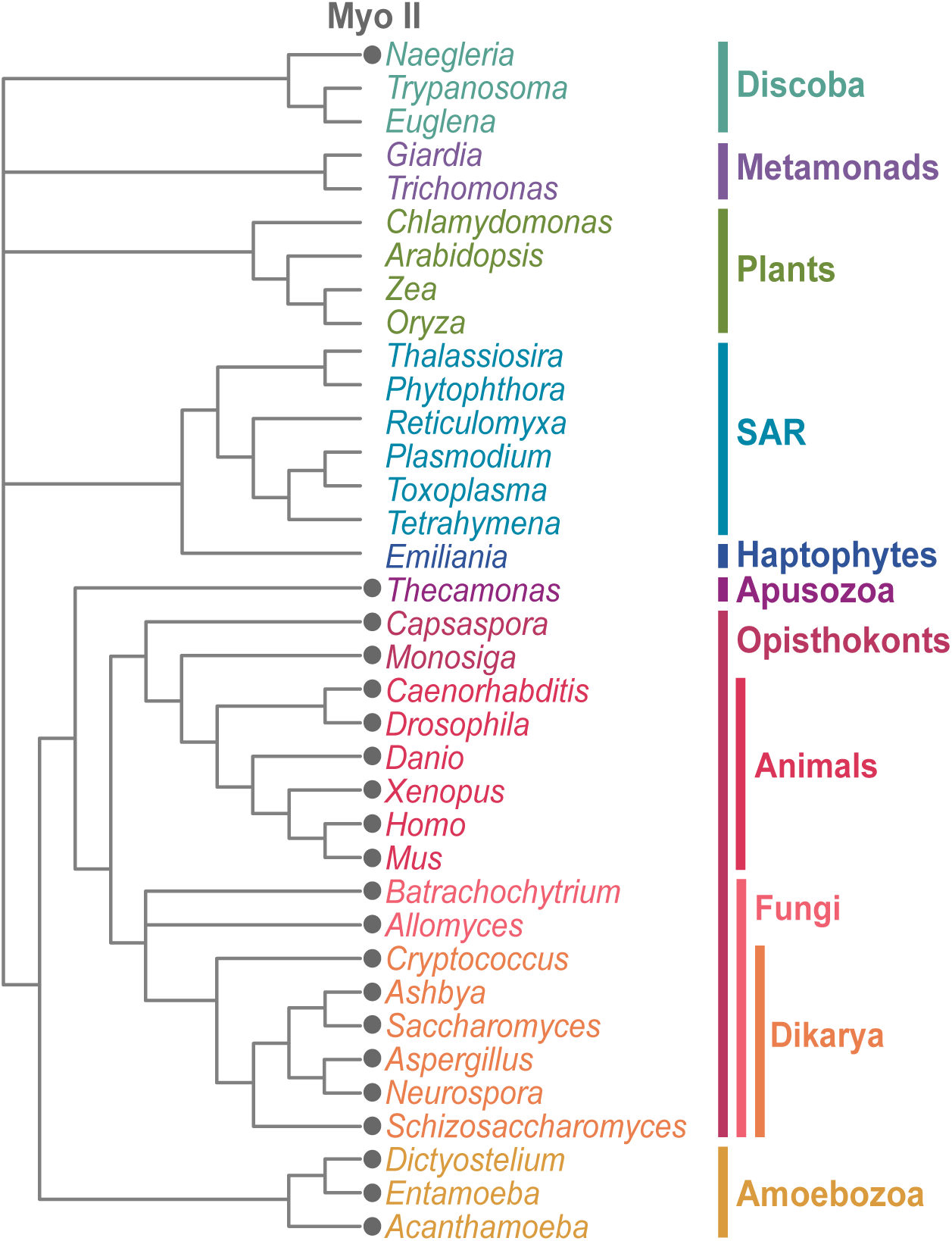
*Naegleria* is evolutionarily distant from well-studied eukaryotic lineages. This diagram illustrates the evolutionary relationships between *Naegleria* and other Eukaryotes. The presence of Myosin II is denoted with a gray circle. This figure has been modified from Velle and Fritz-Laylin, 2019.

While *Naegleria* species spend most of their time as crawling amoebae, under stressful conditions amoebae can differentiate into swimming flagellates. This ∼60 min differentiation involves the assembly of an entire microtubule cytoskeleton from scratch—including transcription and translation of tubulin and its regulators—and has proved to be a valuable model to study microtubules and *de novo* centriole assembly (Fritz-Laylin et al., 2010a; Fritz-Laylin and Cande, 2010; Fritz-Laylin et al., 2016; Fulton and Simpson, 1976; Lai et al., 1979). Although these studies form the basis of most of what we know about *Naegleria*’s cytoskeletal biology, *Naegleria* usually do not have any microtubules at all. The only microtubules found to date in *Naegleria* amoebae are found in the mitotic spindle during closed mitosis (Fulton and Dingle, 1971; Gonzalez-Robles et al., 2009; Walsh, 1984). This means that interphase *Naegleria* amoebae lack microtubules, leading to the assumption that the actin cytoskeleton alone performs all necessary functions for its amoebic lifestyle, particularly cell crawling and phagocytosis.

Despite the fact that actin is considered a pathogenicity factor for the deadly “brain-eating amoeba” *Naegleria fowleri* (Herman et al., 2020; Jamerson et al., 2017; Zysset-Burri et al., 2014), only a handful of *Naegleria* studies over the past five decades have included information on actin (Fritz-Laylin et al., 2010b; Jamerson et al., 2012; Lastovica, 1976; Preston et al., 1990; Sohn et al., 2010; Sohn et al., 2019; Sussman et al., 1984a; Sussman et al., 1984b; Walsh, 2007). Actin has been implicated in *Naegleria*’s extremely rapid (>120 μm/min) crawling motility (King et al., 1983; Preston et al., 1990; Preston and King, 2003), as well as in the formation of *N. fowleri*’s phagocytic cups (Sohn et al., 2010). Analysis of the sequenced genome of the non-pathogenic model system *Naegleria gruberi* provides an important foundation for our understanding of *Naegleria*’s actin biology; the genome encodes dozens of actins and actin regulators, as well as myosin II (Fritz-Laylin et al., 2010b; Herman et al., 2020). The presence of myosin II is interesting, as no other organisms outside of opisthokonts and their close relatives have myosin II (**Fig 1**) (Sebe-Pedros et al., 2014). However, the existing literature lacks a full inventory of *Naegleria*’s actin cytoskeletal regulators in amoebae, quantitative descriptions of the actin structures amoebae build (aside from phagocytic cups), and analyses of the actin nucleation pathways that drive motility and phagocytosis.

Decades of research using model organisms have defined the contributions of actin and microtubule cytoskeletons to opisthokont and amoebozoan motility in molecular detail. Outside of these groups, studies investigating actin-driven cell migration in molecular detail are phylogenetically limited (Frenal et al., 2017; Poulsen et al., 1999). It was therefore unclear if conserved actin-dependent pathways drive motility and phagocytosis in distant eukaryotic lineages. One form of locomotion used by model and non-model organisms which have retained WASP and SCAR/WAVE complex is called “α-motility” (Fritz-Laylin et al., 2017a). This motility relies on highly dynamic, actin-filled pseudopods, and low affinity or non-specific adhesions to the extracellular environment. α-motility is distinct from blebbing motility, where the cell membrane detaches from the underlying actin cortex and blisters outward (Bergert et al., 2012; Yoshida and Soldati, 2006), and from slow mesenchymal motility, in which cells adhere tightly to a two dimensional surface using focal adhesions (Abercrombie, 1980; Gardel et al., 2010; Paluch et al., 2016). Many of these actin-based motility pathways are also invoked for phagocytosis (Cougoule et al., 2004; Heinrich and Lee, 2011; Jaumouille et al., 2019). For example, in *Dictyostelium discoideum*, the SCAR-Arp2/3 pathway used for motility (Davidson et al., 2018; Veltman et al., 2012) is also used for phagocytosis and macropinocytosis (Seastone et al., 2001; Veltman et al., 2016). While actin has clear roles in motility and phagocytosis, the contributions of microtubules to these processes vary greatly by cell type (Etienne-Manneville, 2013). For example, in keratocytes, microtubules are dispensable for motility (Euteneuer and Schliwa, 1984; Nakashima et al., 2015), while in leukocytes they mediate pathfinding and cell shape (Kopf et al., 2020; Renkawitz et al., 2019), and in fibroblasts they are required for directed migration (Liao et al., 1995; Vasiliev et al., 1970). *Entamoeba histolytica* are thought to lack cytoplasmic microtubules ((Meza et al., 2006; Vayssie et al., 2004); but see (Gomez-Conde et al., 2016)), and these amoebae move exclusively by blebbing (Maugis et al., 2010).

Because *E. histolytica* is an amoeba that lacks cytoplasmic microtubules, it could be tempting to speculate that *Naegleria* also moves by blebbing. However, *E. histolytica* is in the amoebozoa lineage, making it more closely related to humans than to *Naegleria*. Moreover, mechanisms of motility vary even among amoebozoans. For example, while *E. histolytica* blebs (Maugis et al., 2010), *D. discoideum* can switch between blebbing and using actin-filled pseudopods for α-motility (Srivastava et al., 2020; Tyson et al., 2014; Zatulovskiy et al., 2014). Therefore, empirically determining the mechanisms that control phenotypes like motility and phagocytosis in an understudied organism is an important step toward understanding its cell biology.

Here, we use genomic analyses to show that *Naegleria gruberi* and the deadly “brain-eating” *Naegleria fowleri* each encode an extensive repertoire of actin cytoskeletal regulators, and these genes are mainly expressed in the amoeboid cell type. We also use fluorescence microscopy to show that *N. gruberi* amoebae build several distinctive actin-rich structures, including Arp2/3-dependent lamellar ruffles. The ability to form these ruffles correlates with increased cell motility and phagocytosis.Additionally, we show that inhibiting contractile, formin-derived networks may impair directional persistence during motility. Collectively, our findings highlight that, despite over a billion years of independent evolutionary history (Betts et al., 2018; Knoll, 2014; Parfrey et al., 2011), *Naegleria* amoebae use actin-based molecular mechanisms strikingly similar to other eukaryotic lineages, supporting an ancient evolutionary origin for α-motility and phagocytosis.

## RESULTS

### *N. gruberi* amoebae lack cytoplasmic microtubules and build diverse actin structures

To define the broad structural differences between the cytoskeletons of *Naegleria*’s two life stages*—* amoebae and flagellates—we examined the actin and microtubule networks of both cell types. We fixed, permeabilized, and stained populations of amoebae and flagellates with phalloidin to label F-actin, and Tubulin Tracker (fluorescently tagged docetaxel) to detect microtubules. Both cell types stained with phalloidin (**Fig 2A**), confirming that an actin cytoskeleton is present in both amoebae and flagellates (Walsh, 2007). Prior studies have also shown that interphase amoebae lack detectable cytoplasmic microtubules (Fritz-Laylin et al., 2010a; Fulton and Dingle, 1971; Walsh, 1984; Walsh, 2007); fitting with this, we did not detect any microtubules in amoebae, while flagellates possessed a pellicle microtubule array in addition to two intensely-stained flagella (**Fig 2A**).

**Fig 2.**
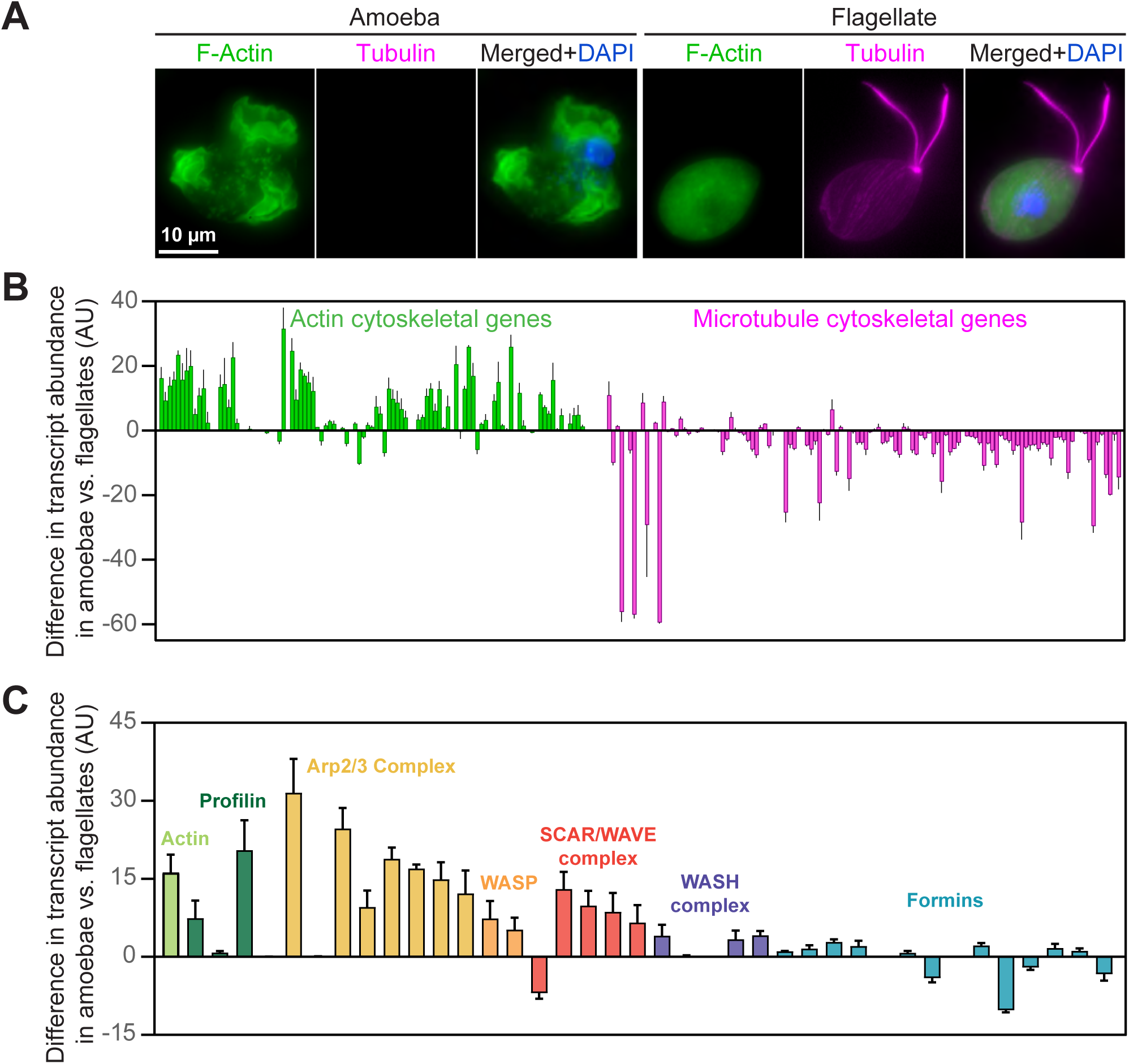
*Naegleria* amoebae employ an actin-based cytoskeletal system, while flagellates use actin and microtubule networks. (**A**) *Naegleria* amoebae are actin-rich but lack cytoplasmic microtubules. A representative NEGM amoeba (left) from a growing population and a flagellate (right) from a population of differentiated (starved) cells were fixed and stained with Alexa Fluor 488 Phalloidin to detect F-actin (green), Deep Red Tubulin Tracker to visualize tubulin (magenta, gamma adjustment of 0.7), and DAPI to label DNA (blue). (**B**) Actin and microtubule cytoskeletal gene expression differs between amoebae and flagellates. Changes in transcript abundance were calculated using gene expression data collected from amoeba or flagellate populations (original data from Fritz-Laylin and Cande, 2010). Data were organized by cytoskeletal system, and display the difference in transcript abundance between the populations in arbitrary units (AU). Positive values indicate higher transcript abundance in amoebae; negative values indicate higher abundance in flagellates. (**C**) *N. gruberi* amoebae induce expression of actin cytoskeletal genes compared to flagellates. A subset of data from panel B was selected to highlight the expression pattern of actin assembly factors. For graphs in B-C, each bar represents the average relative change in transcript abundance between amoebae and flagellates from 3 experiments +/- Standard Deviation (SD).

Actin transcript levels are known to be reduced in *Naegleria* that have transformed into flagellates (Sussman et al., 1984a), although actin protein levels remain constant (Fritz-Laylin et al., 2010a). To expand this work to cytoskeletal regulators, we analyzed actin and microtubule cytoskeletal transcript levels before and after synchronous differentiation from amoebae to flagellates, using the data from our previous investigation of *Naegleria* centriole assembly (Fritz-Laylin and Cande, 2010). Consistent with our previous findings, actin cytoskeletal transcripts (including several of *Naegleria*’s actin paralogs) remained at moderate levels in flagellates (**Fig S1**), although most were more abundant in amoebae (**Fig 2B**). Meanwhile, *Naegleria* amoebae had very low levels of any microtubule cytoskeletal element (**Fig S1A**), while these elements were enriched in flagellates (**Fig 2B**). Taken together with the absence of tubulin staining in amoebae, these data support previous reports that interphase amoebae lack detectable microtubules (Fritz-Laylin et al., 2010a; Fulton and Dingle, 1971; Walsh, 2007), while actin is always present.

We next took a more in depth look at specific actin cytoskeletal genes, many of which are conserved in the “brain-eating amoeba,” *Naegleria fowleri* (**Fig S2**, and (Herman et al., 2020)). Focusing on actin nucleators and nucleation promoting factors, we found that transcript levels for most of these genes were elevated in amoebae relative to flagellates (**Fig 2C**). There are, however, a few notable exceptions; one of the two Arp2 paralogs, one of the two WAVEs, two WASH complex subunits, and six formin family proteins were either expressed at similar levels in both populations, or were preferentially expressed in flagellates (**Fig 2C, Fig S1B**). Because actin is still abundant in flagellates (**Fig 2A-B, Fig S1**), these could be essential components for the flagellate actin cytoskeleton. The abundance of the other transcripts in amoebae indicates that the Arp2/3 complex, its activators (WASP, SCAR/WAVE complex, and WASH complex), and a subset of formins likely facilitate actin-based phenotypes in *N. gruberi* amoebae.

To explore the structures *N. gruberi* builds with this extensive actin machinery, we fixed and stained amoebae to detect F-actin and visualized cytoskeletal morphology using structured illumination microscopy (SIM). This super resolution microscopy revealed a variety of actin structures, including: many 0.2-0.8 μm puncta, 3-6 μm hollow actin spheres, a cell cortex, and thin ruffles (**Fig 3A**). Using widefield fluorescence microscopy to quantify a larger sample size for some of these structures, we found that over 70% of cells had at least 10 actin puncta per cell (**Fig S5I)**, which were present throughout the cell volume—unlike the cortical actin patches used by yeast cells for endocytosis (Adams and Pringle, 1984; Kaksonen et al., 2003; Young et al., 2004). The actin spheres were present in less than a third of cells, and were more difficult to assess without SIM. Because the presence of these structures were variable across controls (**Fig S5I**), we did not conduct further analyses of these structures. The ruffles, however, were clearly present in nearly 70% of cells (**Fig S5I**), and were of particular interest because similar-looking “lamellar” protrusions drive motility in other eukaryotes (Abercrombie, 1980; Abercrombie et al., 1970; Fritz-Laylin et al., 2017a; Fritz-Laylin et al., 2017b; Ingram, 1969). To confirm whether these actin-rich ruffles represented lamellar protrusions, we used scanning electron microscopy (SEM) to visualize the cell surface and observed obvious lamellar protrusions (**Fig 3B**) reminiscent of the sheet-like protrusions and pseudopods of animal cells (Abercrombie et al., 1970; Fritz-Laylin et al., 2017a; Fritz-Laylin et al., 2017b; Ingram, 1969). The presence of these ruffles suggests that *N. gruberi* may move using α-motility, which relies on actin-rich pseudopods, rather than exclusively using blebbing motility like *E. histolytica* (Maugis et al., 2010).

**Fig 3.**
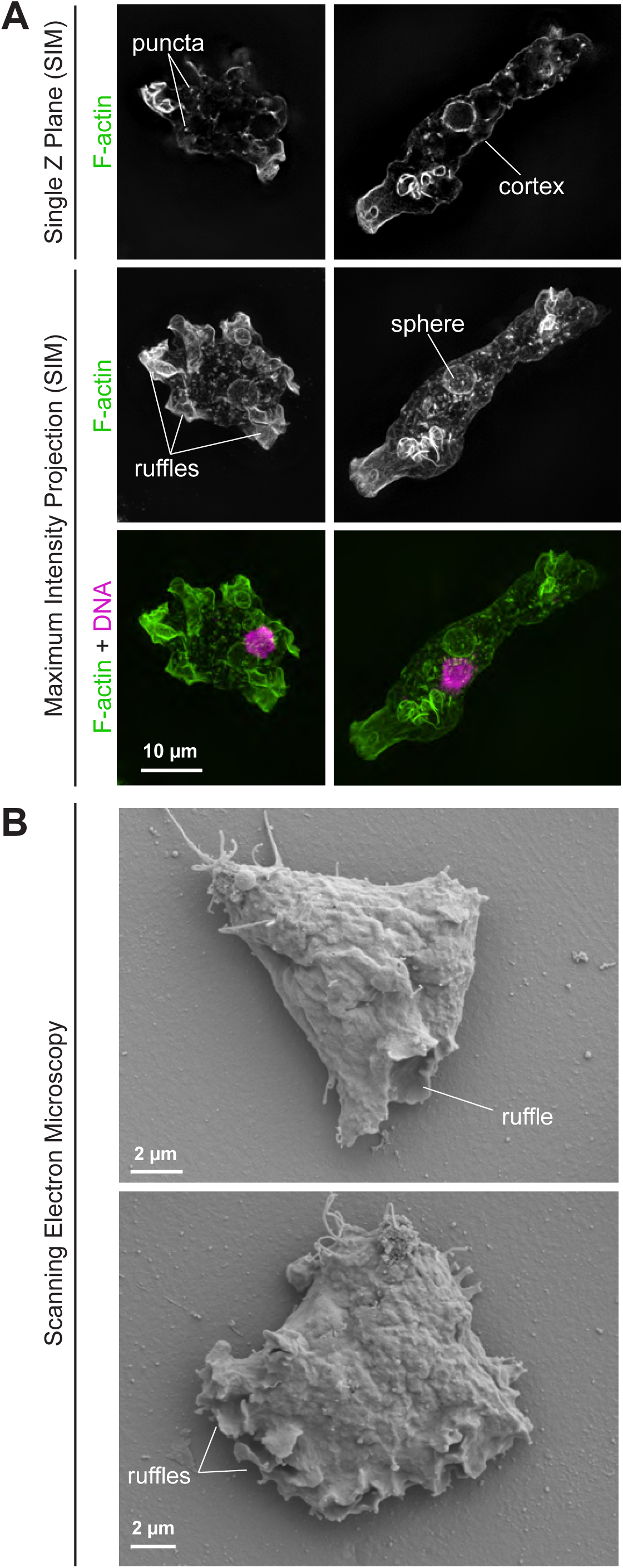
*Naegleria* amoebae build lamellar ruffles and other morphologically distinct structures. (**A**) **Structured illumination microscopy (SIM) of amoebae reveals several morphologically distinct, actin-rich structures.** NEGM amoebae were fixed and stained with Phalloidin to detect actin (green) and DAPI to label DNA (magenta), prior to imaging using SIM to image actin and widefield fluorescence to image DNA. A single Z slice (top) and maximum intensity projections (bottom) are shown for representative control cells (treated with 0.1% DMSO as part of a larger data set shown in Fig 4). Actin structures defined in the text are indicated. **(B) Scanning electron microscopy (SEM) of cells confirms the presence of lamellar ruffles.** NEGM amoebae were fixed and processed for SEM. Ruffles are indicated in 2 representative control cells (treated with 0.1% DMSO).

### Arp2/3 complex activity is required for the assembly of actin-rich puncta and lamellar ruffles

To define how *Naegleria* regulates diverse actin phenotypes, we employed a panel of small molecule inhibitors that target actin and/or actin nucleators (**Fig 4A**). Because *Naegleria* diverged from other eukaryotic lineages >1 billion years ago (Betts et al., 2018; Knoll, 2014; Parfrey et al., 2011), and because these small molecules have not been thoroughly evaluated in *N. gruberi*, we built multiple sequence alignments to assess actin sequence conservation at putative inhibitor binding sites (**Fig S3**). Using the *N. gruberi* actin sequence that has the highest transcript level in amoebae, we found that *Naegleria* actin is 85% identical to mammalian actin, and contains many conserved binding sites for the small molecule inhibitors described below (**Fig S4**). An *N. fowleri* actin sequence nearly identical to previous work (Sohn et al., 2010) has an amino acid sequence 99% identical to that of *N. gruberi*, and contains identical residues at drug binding sites (**Fig S3-S4**), suggesting that drugs effective against *N. gruberi* actin may also work in other *Naegleria* species.

**Fig 4.**
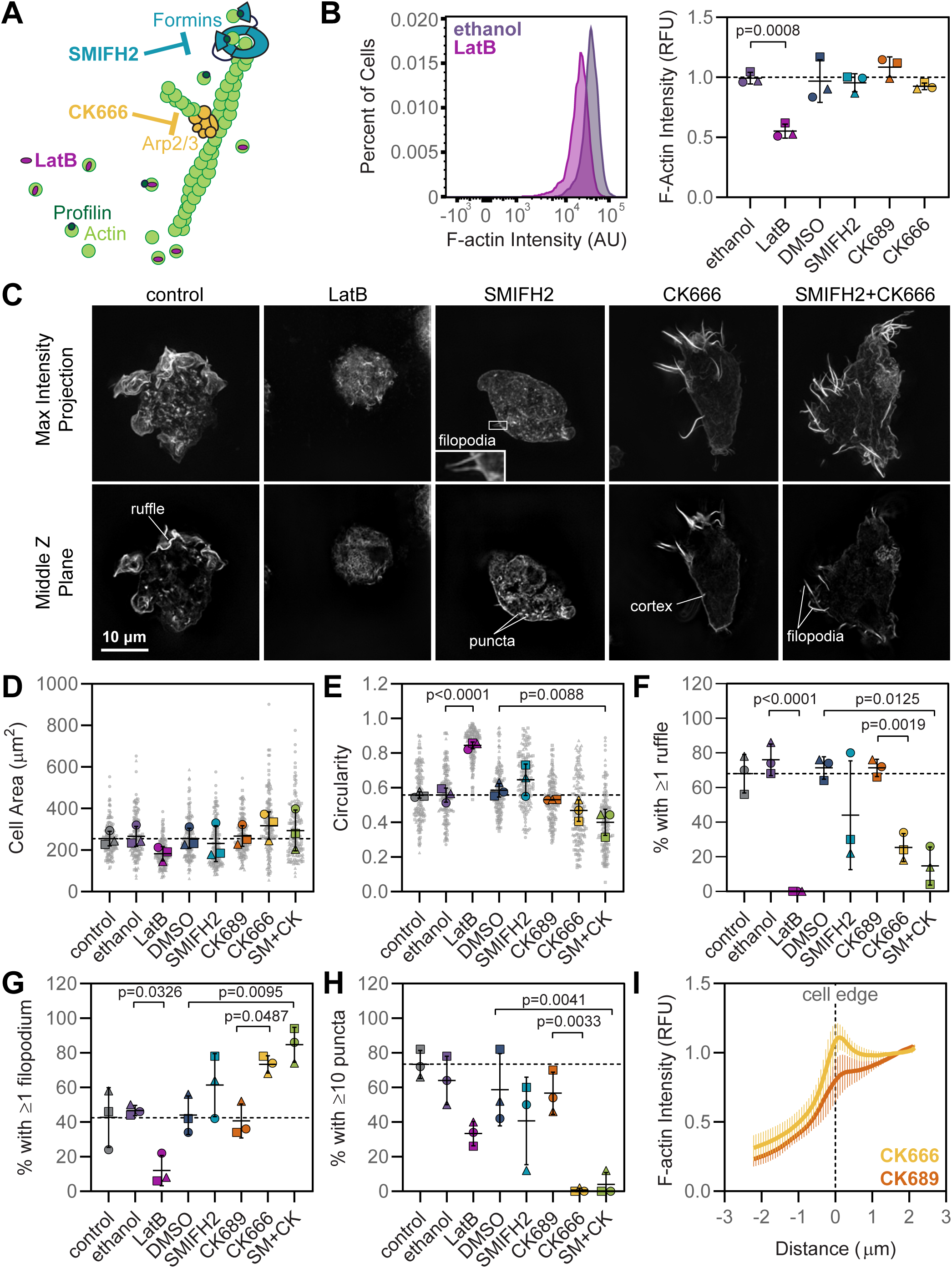
Small molecule inhibitors of the actin cytoskeleton alter *Naegleria* morphology. **(A) Small molecule inhibitors were implemented to alter *Naegleria* actin dynamics.** Distinct actin assembly pathways are blocked by small molecule inhibitors. Drugs are matched with vehicle or inactive controls as follows: LatB/ethanol, SMIFH2/DMSO, CK-666/CK-689. **(B) LatB decreases global actin polymer levels measured by flow cytometry.** NEGM cells were incubated in media +/- inhibitors or specific controls for 10 min, then fixed and stained with phalloidin. Total fluorescence was then measured using flow cytometry to analyze 20,000 cells for each sample. One representative histogram (left) is shown from a single replicate comparing LatB treatment to its vehicle control. See Figure S4 for additional histograms. Average F-actin intensities (right) were normalized to the stained control for all conditions. Each point represents the average normalized fluorescence intensity of F-actin staining from one experimental replicate. **(C) Fluorescence microscopy shows morphological differences following inhibitor treatments.** NEGM cells were treated with inhibitors and controls as in B, but were fixed and stained for microscopy using DAPI to label DNA, and phalloidin to detect F-actin. Cells were analyzed using structured illumination microscopy (this figure) and widefield fluorescence microscopy (see Fig S3). Maximum intensity projections and single Z planes from SIM are shown for representative cells, and filopodia on a SMIFH2-treated cell are magnified 4x in an inset. **(D-E) Actin inhibitors alter cell shape, but not size.** Widefield fluorescence microscopy data from experiments shown in C were used to analyze 50 cells per condition for each of three independent replicates (150 cells total/treatment). Outlines were drawn around cells in Fiji to measure (D) cell area, and (E) circularity. Small gray symbols represent individual cells, while larger symbols represent the average of each of three experimental replicates (squares, circles, and triangles). **(F-H) Treatment with actin inhibitors alters cell morphology.** The same cells analyzed in D-E were also classified by their morphological features. Cells were scored blindly to quantify the percent of cells that possessed: (F) at least one ruffle, (G) at least one filopodium, and at least ten puncta. Symbols represent the percentages from three experimental replicates. For graphs in B and D-H, solid lines represent the means calculated from three experimental replicates, +/- SD, and statistical significance was determined using an ordinary ANOVA followed by Tukey’s multiple comparison test. Dashed lines indicate the control value. **(I) CK-666 treatment enhances cortical actin levels.** Pixel intensity linescans (line width = 50 px, representing 3.25 μm) for F-actin staining were drawn in a single Z plane bisecting the cell edge, and values were normalized to the average intensity 1.5-2.0 μm into the cell, which was set to 1. Curves represent the average relative intensity +/- SD for 3 experimental replicates, each encompassing 5 cells (see Fig S5 for representative images and all individual line scans).

To inhibit actin monomer/polymer dynamics, we used latrunculin B (LatB), which prevents actin assembly (Spector et al., 1983), as well as jasplakinolide (Bubb et al., 1994) or phalloidin (Lengsfeld et al., 1974) to prevent filament turnover, and cytochalasin D to prevent elongation (Flanagan and Lin, 1980). To impair actin nucleation pathways, we used CK-666 to inhibit Arp2/3 complex activity (Hetrick et al., 2013; Nolen et al., 2009), and SMIFH2 to broadly inhibit formin activity (Rizvi et al., 2009). In addition to inhibiting formins, SMIFH2 was also recently shown to inhibit some myosins (Sellers et al., 2020), and therefore may be better described as an inhibitor of contractile networks. Regardless, targeting formins with SMIFH2 has been a reliable indicator for the involvement of formins in other systems (Fattouh et al., 2015; Rana et al., 2018; Velle and Campellone, 2018).

To determine if these small molecule inhibitors impact specific actin networks or globally disrupt actin polymerization, we assessed their effects on total actin polymer content. We fixed cells after 10 min of exposure to small molecules or controls, stained polymerized actin using fluorescently labeled phalloidin, and analyzed total cell fluorescence using flow cytometry (Kakley et al., 2018). We compared the histograms of F-actin intensities for small molecules to their respective controls (LatB vs. ethanol vehicle control; jasplakinolide, phalloidin, cytochalasin D, or SMIFH2 vs. DMSO vehicle control; or CK-666 vs. CK-689 inactive control), and normalized the fluorescence intensities for each condition to the stained control sample from that replicate (**Fig 4B, Fig S5B-H**). Only LatB treatment resulted in a peak shifted towards zero, indicative of lower levels of actin polymer. On average, LatB reduced total cellular F-actin content by 44.5% (**Fig 4B**), consistent with its effects as an actin depolymerizing agent (Spector et al., 1983). Cells treated with jasplakinolide, phalloidin, cytochalasin D, SMIFH2, or CK-666 all had similar levels of actin polymer to controls. Because cells treated with unlabeled phalloidin as part of the experiment stained just as well as control cells with labeled phalloidin following fixation and permeabilization (**Fig S5F**), it seems likely that the phalloidin either never reached the cytoplasm in live cells, or was efficiently cleared from the cell by efflux pumps. Collectively, these results show that while LatB decreases the cellular actin polymer content, the application of the other small molecules used in this study do not result in gross defects in F-actin abundance at these concentrations.

We next assessed whether specific inhibitors alter the morphology of individual *Naegleria* amoebae. We again treated cells for 10 minutes with inhibitors or controls, fixed and stained cells to detect F-actin, and analyzed cells using widefield and super resolution microscopy (**Fig 4C, Fig S5A**). SIM imaging revealed that control-treated cells displayed the previously identified actin structures, including ruffles and puncta (**Fig 3A**). Cells treated with LatB were rounded and lacked any distinguishing features (**Fig 4C**), a result which fits with the global defects in actin polymerization measured by flow cytometry (**Fig 4B**). While treatment with SMIFH2 did not reliably correlate with any morphological phenotypes we analyzed, CK-666-treated cells (and cells treated with both CK-666 and SMIFH2) had a robust actin cortex, and striking actin-rich spikes, reminiscent of filopodia (**Fig 4C**). While control cells and SMIFH2-treated cells also had these filopodia-like protrusions, they were often dimmer and shorter (**Fig 4C**, inset, **Fig S5I**).

To quantify the prevalence of these morphological features in control and inhibitor-treated cell populations, we imaged 150 cells per treatment using widefield fluorescence microscopy, and quantified the cell size (2D area), shape, and presence of actin structures (**Fig 4D-H**). While cell size did not differ significantly between treatments (**Fig 4D**), cells were more circular (with values closer to 1) following LatB treatment and more elongated (with values closer to 0) following combined treatment with CK-666 and SMIFH2 (**Fig 4E**). Arp2/3 complex inhibition significantly decreased cell ruffles (**Fig 4F**), while increasing the percent of cells with filopodia-like structures (**Fig 4G**). Further, actin puncta were very rare in CK-666 treated cells; less than 1% of CK666-treated cells had at least ten 0.2-0.8 μm actin-rich foci (**Fig 4H**), suggesting these structures are dependent on Arp2/3 complex activity. Finally, the cell cortex was noticeably more intense in CK-666 treated cells (**Fig 4C**). To quantify this observation, we drew line scans centered on the cell edge (**Fig 4I, Fig S6**). Compared to the inactive control CK-689, CK-666 treatment resulted in a local maximum in fluorescence intensity at the cell edge. Collectively, these results suggest that the Arp2/3 complex is important for forming ruffles and actin puncta, and in the absence of Arp2/3 activity, cells preferentially build filopodia-like structures and may allocate more actin to the cell cortex.

### Inhibiting actin nucleation by the Arp2/3 complex reduces *Naegleria* cell speed while inhibiting formins impairs directional persistence

To assess whether the morphological changes observed with the actin inhibitors translated to defects in cell migration, we introduced inhibitors during live imaging of crawling cells. We then directly compared pre- and post-treatment speeds of individual cells by tracking cells 5 min before, during, and 5 min after treatment (**Fig 5A-B, Movie S1**). To visualize this data, we plotted the pre-treatment vs post-treatment cell speeds with respect to a line with a slope of 1, which indicates no change in speed. Cells that fall under the line moved slower after treatment, while cells above the line moved faster. The speed of control cells changed an average 1.3+/-1.6 µm/min (**Fig 5C**), which is a measure of the variability in this system. Similar to control treatment, exposure to ethanol, DMSO, or CK-689 controls, or to jasplakinolide, phalloidin, or cytochalasin D, did not reduce cell speeds at the concentrations tested (**Fig 5C, Fig S7A**). In contrast, 100% of cells treated with LatB exhibited slower cell speeds (29.8+/-4.3 µm/min pre-treatment vs. 4.6+/-2.3 µm/min post-treatment). This implies that actin polymerization is critical for the motility of *Naegleria* amoebae.

**Fig 5.**
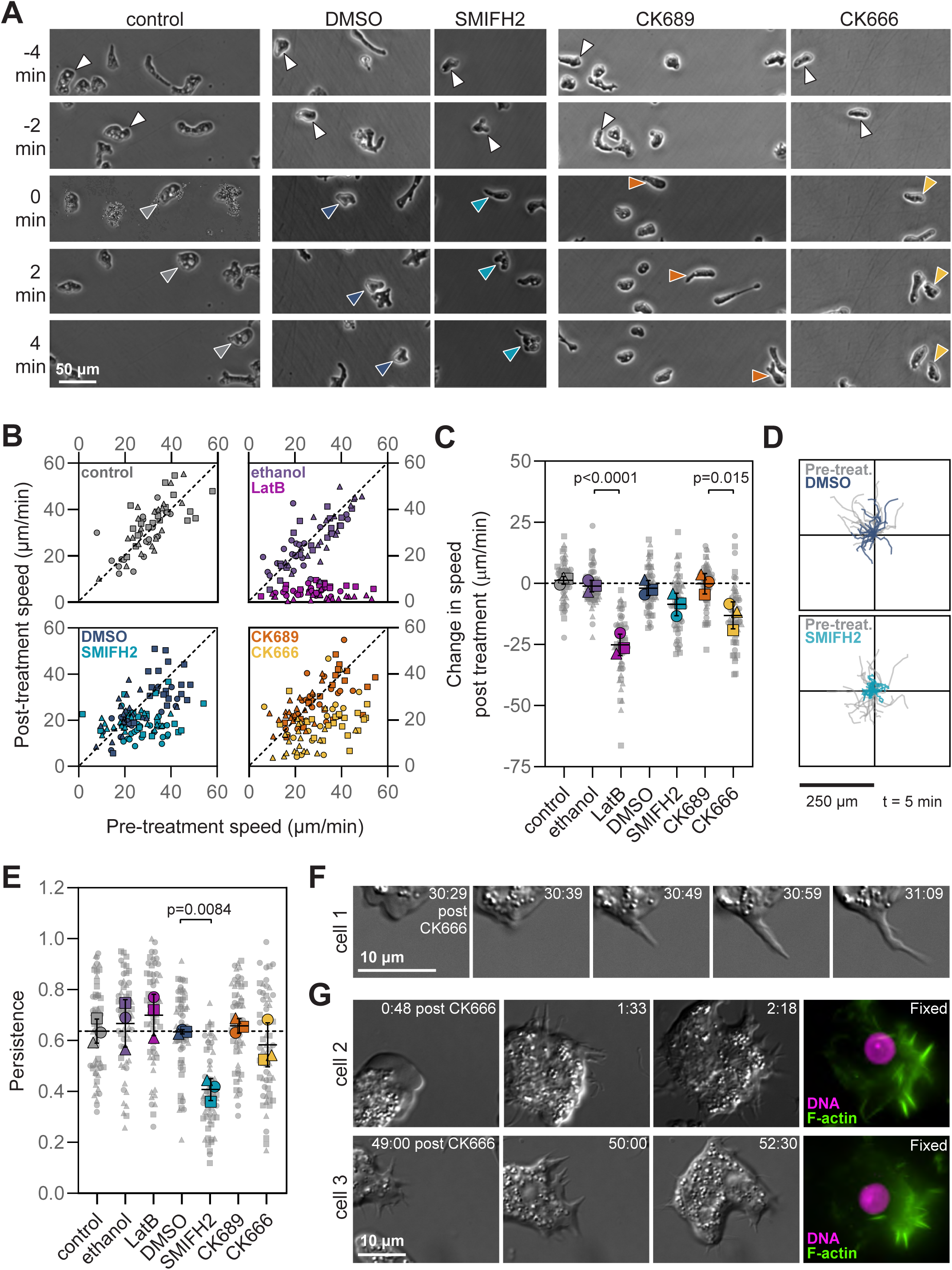
Inhibition of actin nucleation pathways impairs *Naegleria* cell crawling. **(A) Live imaging cells during inhibitor treatments reveal defects in motility.** Crawling amoebae were imaged using phase/contrast microscopy for 5 min. After 5 min (at t=0 min), an equal volume of each of the indicated small molecules or controls were added (diluted in buffer; “control” indicates buffer alone was added), and imaging proceeded for 5 min. Arrowheads indicate the position of a representative cell over time. **(B) All LatB-treated cells and most CK-666-treated cells move slower after treatment.** Cells were treated as in A. 20 randomly selected cells from the center of the field of view at t=0 were tracked to calculate average speeds before and after treatment. Each point represents the speed of a cell before treatment (plotted on the x axis) and after treatment (plotted on the y axis). The dashed line represents no change. 3 experimental replicates are indicated using different shapes (squares, triangles, and circles). Two data points are not displayed, as they are off the limit of the x axis (LatB (79.3, 12.9), and CK-666 (60.8, 23.6)). **(C) LatB and CK-666 decrease average cell speed.** The data collected panel B were used to calculate the change in cell speed post-treatment. Each smaller gray symbol represents a single cell, while larger symbols represent the averages from experimental replicates, coordinated by shape (n=3, 20 cells per trial, 60 cells total per condition). The dashed line is set to zero. **(D) SMIFH2-treated cells explore less area than control cells.** Cell migration data from B were plotted such that for tracks prior to treatment (gray traces), the -5 min time point was centered at the origin, while for tracks after treatment (shown in color) the t=0 time point was centered at the origin. Tracks from 20 cells from one representative experiment are shown. **(E) SMIFH2 treatment impairs directional persistence.** Directional persistence was calculated for each post-treatment cell tracked in B by dividing the maximum displacement from the start of the track by the total path length. Each small gray symbol represents the persistence of a single cell, while larger symbols represent experimental averages. For graphs in C and E, black lines indicate the mean +/- SD calculated from 3 experimental replicates, and statistical significance was determined using an ordinary ANOVA followed by Tukey’s multiple comparison test. **(F) Filopodia-like protrusions grow de novo following CK-666 addition.** Cells were imaged using DIC microscopy at high magnification, and treated with CK-666 during imaging. Panels show time lapse images of a representative cell generating a filopodium-like protrusion approximately 30 min after treatment. **(G) Filopodia-like protrusions are actin-rich.** Two additional cells treated and imaged as in (F) were fixed and stained on the microscope after 2 min and 18 sec of treatment (top, cell 2) or 53 minutes of treatment (bottom, cell 3). Cells were simultaneously fixed, permeabilized, and stained, with DAPI to detect DNA (magenta), and Phalloidin to detect F-actin (green). Times post treatment for F-G are shown in min:s.

Having determined that actin polymerization is key to *Naegleria* motility, we next sought to determine which actin nucleation pathways drive this behavior. To assess the contributions of the Arp2/3 complex, we treated cells with CK-666. CK-666 treatment reduced speed in 90% of cells, and on average cells moved 13.1+/-5.5 µm/min slower after CK-666 treatment (**Fig 5 B-C**). This impaired motility following CK-666 treatment suggests that the Arp2/3 complex is a major contributor to *Naegleria*’s actin-mediated motility.

While CK-666 reduced cell migration, treatment with SMIFH2 did not result in a statistical difference in cell migration speeds. There were, however, obvious differences in the way SMIFH2-treated cells moved; cells often contacted nearby cells and then moved in circular patterns (**Movie S1, Fig 5A**). To visualize these behaviors, we used worm plots to display the tracking data for 20 cells collected from 5 min before and 5 min after treatment. For tracks prior to treatment (gray traces), the -5 min time point was centered at the origin, while the t=0 time point was centered at the origin for post-treatment tracks (colored traces). From these plots, it is clear that SMIFH2-treated cells do not explore the same amount of surface area as untreated or DMSO-treated cells (**Fig 5D**). As a more formal test of this, we compared the maximum displacement values, which measure the furthest a cell reaches from the starting point over the course of cell tracking; SMIFH2-treated cells had an average maximum displacement of 36.2+/-4.6 μm after treatment, compared to 95.2+/-15.6 μm before treatment. Further, we calculated the directional persistence by dividing the maximum displacement by the total path length (the most direct route would have a value of 1). DMSO-treated cells had a persistence of 0.63+/-0.04, while treatment with SMIFH2 significantly decreased the directional persistence to 0.41+/-0.04 (**Fig 5E**). Collectively, these data indicate that actin-based motility in *N. gruberi* is largely dependent on actin nucleation by the Arp2/3 complex, while directional persistence during cell migration may rely on formin-nucleated actin networks.

Actin nucleators have been shown to collaborate in the production of a variety of cell phenotypes, including lamellipodial protrusions (Bovellan et al., 2014; Isogai et al., 2015; Kage et al., 2017; Velle and Campellone, 2018; Yang et al., 2007). To address the possibility that the Arp2/3 complex and formins could either cooperate or compensate for each other to drive motility, we set out to determine if inhibiting both would exaggerate the observed motility defects. Cells treated with both CK-666 and SMIFH2 combined were neither significantly slower nor less directionally persistent than those treated with CK-666 or SMIFH2 alone (**Fig S7B-D**). While these data do not exclude the possibility for some amount of nucleator crosstalk, they are not consistent with a requirement for formin and Arp2/3 complex synergy in *Naegleria*’s actin-dependent cell migration.

Because some systems do not strictly require the Arp2/3 complex for motility (Rotty et al., 2017), we looked more closely at the residual motility of CK-666-treated cells. While a fraction of cells continued to move after treatment using a mechanism reminiscent of blebbing (**Movie S1**), a larger percentage of cells developed filopodia-like protrusions, similar to the actin-enriched structures we previously observed (**Fig 4C**). Higher resolution imaging of these CK-666-treated cells revealed that these structures form *de novo* from the leading edge of the cell and grow outward at an average maximum rate of 0.90+/-0.47 μm/s (maximum growth rate for actin-enriched filopodia over a 3 s window, averaged from 4 cells) (**Fig 5F**). We also imaged crawling cells before and during treatment with CK-666, and fixed and stained these cells for F-actin during the time course (**Fig 5G, Movie S2**). This confirmed that these structures are actin-rich, and therefore are likely the same structures we previously observed in bulk fixed cells (**Fig 4C**). Collectively, these data show that although *Naegleria*’s main form of cell migration relies on Arp2/3-dependent lamellar protrusions, cells can still migrate, to a degree, under Arp2/3 inhibition.

### Phagocytosis requires actin assembly, and is enhanced by Arp2/3 complex activity

In diverse cell types including *Dictyostelium* amoebae and mammalian leukocytes, the actin networks that promote the assembly of lamellar pseudopods also promote phagocytosis (Davidson et al., 2018; Jaumouille et al., 2019; Seastone et al., 2001; Veltman et al., 2012; Veltman et al., 2016). To test the hypothesis that Arp2/3-dependent actin networks drive phagocytosis as well as cell motility in *Naegleria*, we treated cells with LatB or CK-666 and incubated cells with GFP-expressing bacteria. After fixing cells and visualizing GFP-bacteria by microscopy, we observed that many control cells (untreated, ethanol, and CK-689 treated) were associated with robust GFP signal, while CK-666 treated cells had considerably less GFP intensity, and LatB-treated cells were associated with little-to-no fluorescence (**Fig 6A**).

**Fig 6.**
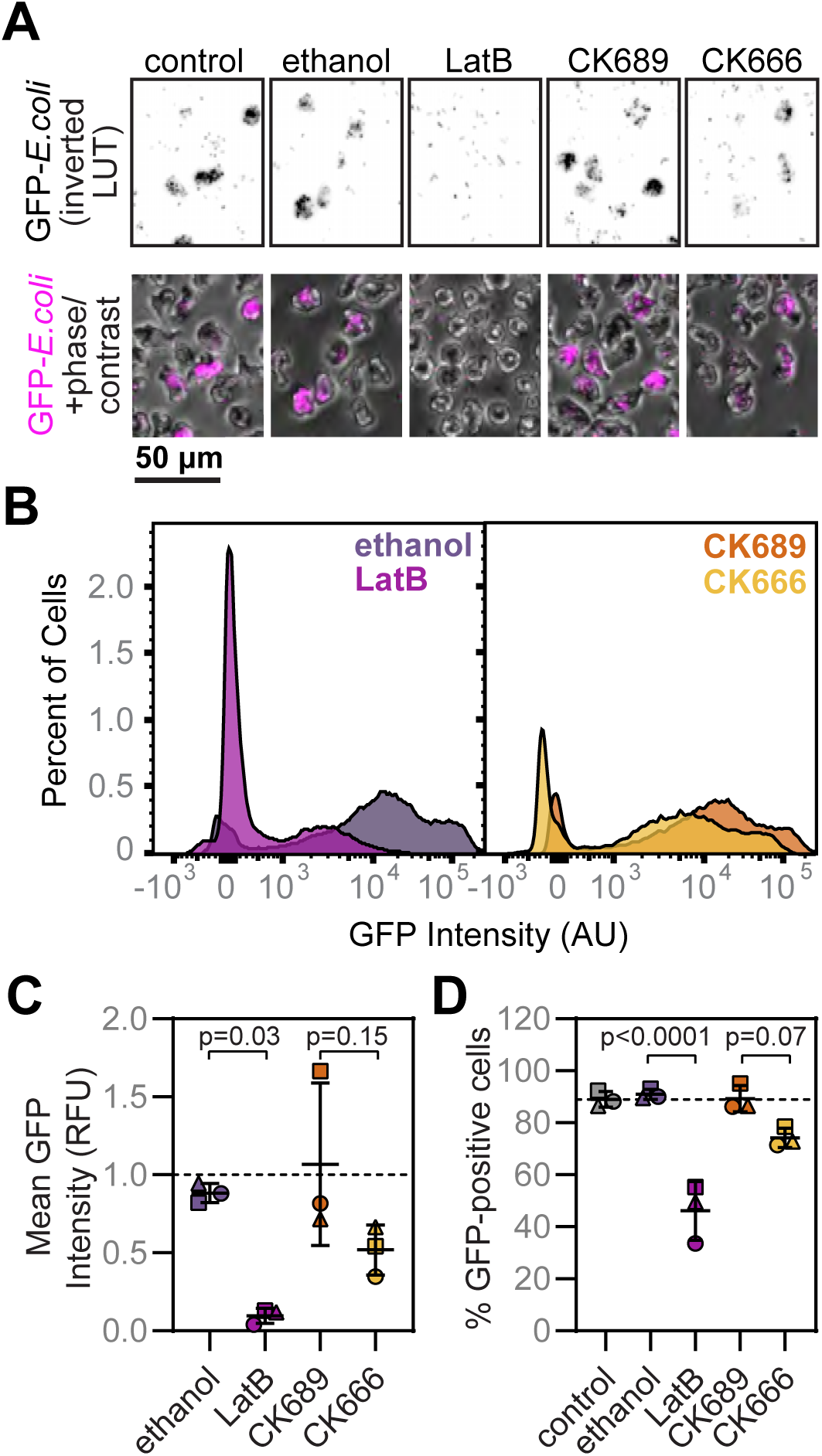
Phagocytosis is less efficient following Arp2/3 complex inhibition. **(A) LatB and CK-666 reduce bacteria-associated fluorescence of amoebae visualized by microscopy.** NEGM amoebae were starved for 1 h, then incubated with controls or inhibitors and GFP-expressing *E. coli* for 45 min. Cells were then fixed and visualized at low magnification. GFP intensity is shown using an inverted LUT (top), and in magenta (bottom). **(B) LatB or CK-666 treatments reduce bacteria-associated GFP signal measured by flow cytometry.** Cells were starved, treated, and fixed as in A. Then, the GFP intensities of 30,000 cells per condition were measured by flow cytometry. Representative histograms display the percent of cells (after gating to select only intact, single cells) plotted against GFP intensities for ethanol and LatB treated cells (left), or CK-689 and CK-666 treated cells (right). Plots are from one of three experimental replicates (see Figure S8 for all histograms). **(C) LatB-treatment reduces average bacteria-associated fluorescence.** For each experimental replicate (square, circle, triangle), the average fluorescence intensity of control and treated cells were normalized to the average intensity of a buffer-only control. Each point represents the average normalized fluorescence intensity of intact, single cells. **(D) Fewer LatB-treated cells are associated with bacteria than control cells.** The population of buffer-only control cells was used to gate a GFP-positive population for each experimental replicate. Each point represents the percent of treated cells falling within the GFP-positive gate. For graphs in C-D, lines indicate the mean of three experimental replicates +/- SD. Statistical significance was determined using an ordinary ANOVA followed by Tukey’s multiple comparison test.

To quantitate this observation, we measured the levels of cell-associated bacteria using flow cytometry. Compared to the ethanol vehicle control, the peak in intensity for GFP-positive LatB-treated cells was shorter and shifted left (towards 0), and the vast majority of cells had virtually no GFP fluorescence (**Fig 6B**). While not as dramatic, CK-666-treated cells also had a shorter, left shifted peak, and a higher peak representing GFP-negative cells than CK-689-treated cells (**Fig 6B, Fig S8A**). For LatB treatment, these shifts translated to a significantly lower mean intensity compared to ethanol alone, and a 50.7+/- 11.9% reduction in the percent of positive cells (**Fig 6C-D**), indicating that phagocytosis does indeed involve actin polymerization. For cells treated with CK-666, neither the change in GFP intensity, nor the percentage of positive cells reached our threshold for statistical significance. However, the three biological replicates had the same trend, suggestive of a defect in phagocytosis despite variability among individual cells (**Fig S8A**).

To gain more insight into the nature of the residual “bacteria-positive” LatB and CK-666-treated cells, we repeated the experiment, and stained cells using fluorescently labeled phalloidin to visualize the actin cytoskeleton. Cells had two distinct types of GFP staining: intense staining that appeared to correspond to external, intact bacteria, as well as weaker staining that appeared to represent internalized bacterial remnants (**Fig S8B**). While both types of staining were often observed in control cells, LatB and CK-666 treated cells were typically found with only extracellular bacteria. This suggests that the “GFP positive” LatB and CK-666 cells identified by flow cytometry may have been associated with extracellular bacteria, and therefore may not represent *bona fide* phagocytic events. More in depth studies are required to decipher if defects associated with Arp2/3 inhibition exist in binding bacteria, internalizing bacteria, digesting bacteria, or all three.

## DISCUSSION

Our work demonstrates that *Naegleria* uses sophisticated actin cytoskeletal machinery, including the Arp2/3 complex, three Arp2/3 activators, and formin family proteins. These proteins are primarily expressed in the microtubule-free, amoeboid cell state. Using small molecule inhibitors, we show that cells build Arp2/3-dependent actin puncta and lamellar protrusions, and assemble filopodia and cortical actin networks during Arp2/3 inhibition. We also show that Arp2/3 inhibition impairs cell motility and phagocytosis, while inhibiting formin-derived contractile networks with SMIFH2 impairs directional persistence. Because Arp2/3-derived networks also promote lamellar protrusions, α-motility, and phagocytosis in opisthokonts and amoebozoans (Davidson et al., 2018; Jaumouille et al., 2019; Seastone et al., 2001; Veltman et al., 2012; Veltman et al., 2016), our results suggest that the Arp2/3-dependent mechanisms that drive motility and phagocytosis are evolutionarily ancient, and can function independently of a microtubule cytoskeleton.

The complexity of *Naegleria* actin networks is in stark contrast to some early-diverging eukaryotes that have highly simplified actin cytoskeletons. For example, the intestinal parasite *Giardia lamblia* has an extremely divergent actin, and although its expression is important for endocytosis, membrane trafficking, and cytokinesis (Paredez et al., 2011; Paredez et al., 2014), *Giardia* lacks homologs of nearly every known actin binding protein (Morrison et al., 2007). *Trichomonas vaginalis* is a metamonad relative of *Giardia*. Like *Naegleria, T. vaginalis* can exist as an amoeba or a flagellate. The *T. vaginalis* amoebae generate actin and fimbrin rich protrusions, and the genome encodes several actin binding proteins, despite lacking myosins and some actin bundling proteins (Carlton et al., 2007; Kusdian et al., 2013). Together, these data suggest that *Giardia*’s lack of actin binding proteins is a result of divergence, and raises the possibility that the ancestor of the metamonads, heteroloboseans, amoebozoans, and opisthokonts likely had a veritable smorgasbord of actin regulators and may have been able to undergo crawling motility (Fritz-Laylin et al., 2017a; Fritz-Laylin et al., 2010b; Sebe-Pedros et al., 2013; Sebe-Pedros et al., 2014; Velle and Fritz-Laylin, 2019).

*Naegleria*’s repertoire of actin cytoskeletal genes includes ≥30 actins, 78 total actins and Arps (Fritz-Laylin et al., 2010b), as well as two Arp2s and two SCAR/WAVEs. These numbers are similar to those of *Dictyostelium discoideum*, whose genome encodes 17 identical actins, and 41 total actins and Arps (Joseph et al., 2008). This abundance of actin genes likely indicates the importance of actin to an amoeboid lifestyle. Moreover, in environments where other organisms produce actin toxins, it is possible some of these additional genes encode toxin-resistant actins, similar to the strategy used by *Chlamydomonas reinhardtii* (Onishi et al., 2016). *Naegleria’s* two Arp2 genes indicate the potential to form multiple types of Arp2/3 complexes, which could be optimized for different tasks like in mammalian cells (Molinie et al., 2019). Our analysis does not support robust expression for one of these Arp2s in amoebae or flagellates, so a significant role for multiple Arp2/3 complexes seems unlikely for these life stages. On the other hand, *N. gruberi*’s two SCAR/WAVE genes are differentially expressed, suggesting different SCAR/WAVE complexes exist in amoebae and flagellates. Mammalian cells use multiple WAVEs (which diversified within the animal lineage (Kollmar et al., 2012; Veltman and Insall, 2010) to form different types of ruffles (Suetsugu et al., 2003). Further work is needed to determine if these distinct SCAR/WAVE complexes have different cellular functions and Arp2/3 activation potencies in *Naegleria*.

Because *Naegleria* amoebae lack cytoplasmic microtubules (Fritz-Laylin et al., 2010a; Fulton and Dingle, 1971; Walsh, 1984), our work here demonstrates that motility and phagocytosis can function independent of a microtubule cytoskeleton in a natural context. In other cell types, microtubules are important for cell migration across 2D surfaces (Liao et al., 1995; Vasiliev et al., 1970), maintaining polarity (Zhang et al., 2014), pathfinding in complex environments (Renkawitz et al., 2019), and cell shape (Kopf et al., 2020). Microtubules have also been found to contribute to phagocytosis in mammalian cells, particularly during CR3-mediated phagocytosis (Allen and Aderem, 1996; Lewkowicz et al., 2008; Newman et al., 1991). Aside from *Naegleria*, the only other natural system thought to lack cytoplasmic microtubules, to our knowledge, is *Entamoeba histolytica* ((Meza et al., 2006; Vayssie et al., 2004); but see (Gomez-Conde et al., 2016)), which move exclusively by blebbing motility (Maugis et al., 2010). Our live imaging data show that while some *Naegleria* cells can generate rapid bleb-like protrusions to move, they typically use phase dense, lamellar protrusions (**Movie S1-S2**). Taken together with our SEM images and F-actin staining data, these protrusions are lamellar and actin-rich, therefore suggesting that unlike *E. histolytica, N. gruberi* amoebae use α-motility.

One of the most extensively studied types of motility is the 2D migration of adherent epithelial and mesenchymal cells (Gardel et al., 2010). While this type of motility is at least an order of magnitude slower than α-motility (Fritz-Laylin et al., 2017a), it does share some key features; Arp2/3-activating factors (particularly the SCAR/WAVE complex) promote branched actin networks (Fritz-Laylin et al., 2017a; Fritz-Laylin et al., 2017b; Machesky and Insall, 1998; Mullins et al., 1998; Svitkina and Borisy, 1999), these networks push against the cell membrane (Mullins et al., 2018), and the protrusions formed by this process are lamellar (Abercrombie, 1980), even when crawling through a 3D environment (Fritz-Laylin et al., 2017b). While *Naegleria* was assumed to be using α-motility, our work adds important verification for this, as we confirmed the presence of actin rich lamellar protrusions through fluorescence and electron microscopy.

We also show that Arp2/3-dependent lamellar protrusions correlate with cell motility, as treating cells with the Arp2/3 inhibitor CK-666 impaired both their assembly and cell motility. Further, Arp2/3 inhibition impaired phagocytosis, suggesting that like other organisms, there is some overlap in the pathways responsible for moving and eating. While inhibiting formins with SMIFH2 often resulted in more subtle phenotypes than Arp2/3 inhibition, there was an obvious defect in cell motility. SMIFH2-treated cells did not significantly slow down, but lost directionality and were unable to maintain a polarized leading edge. Because SMIFH2 is capable of inhibiting myosins in addition to formins (Sellers et al., 2020), either effect could disrupt contractile networks that maintain membrane tension, resulting in the cells losing their ability to maintain a polarized front and rear (Hind et al., 2016; Houk et al., 2012). The precise mechanisms regulating cell migration and polarity in *Naegleria*, therefore, remain to be elucidated. Overall, our data indicate that lamellar protrusions built by Arp2/3 complex drive *Naegleria* motility, suggesting α-motility is widespread and potentially evolutionarily ancient.

Filopodia are also widespread throughout the tree of life, and are thought to have evolved during early eukaryotic evolution (Sebe-Pedros et al., 2013). Several cell types naturally possess filopodia to serve a variety of functions including motility (Jacquemet et al., 2015; Meyen et al., 2015). Mammalian leukocytes treated with CK-666 or lacking Arp2/3 complex show an enrichment in filopodial protrusions (Fritz-Laylin et al., 2017b; Rotty et al., 2017). We observed similar enrichment in filopodia upon CK-666 treatment. While it seems reasonable that when cells cannot form ruffles they could switch to a different type of protrusion using another nucleator, we also observed these filopodia in cells treated with both CK-666 and SMIFH2. If these filopodia are nucleated by formins, it is possible that SMIFH2 does not inhibit all 14 of *Naegleria*’s formins, or perhaps *Naegleria* may have additional actin nucleators not found in opisthokonts. In any event, the increase in filopodia upon CK-666 treatment highlights another similarity in actin networks from organisms across the eukaryotic tree.

Collectively, our work lays a foundation for studying actin cytoskeletal biology in *Naegleria*. This work also has implications for the “brain-eating amoeba” *N. fowleri*. The lack of effective, reproducible treatment options for *N. fowleri* infections has resulted in a >90% case fatality rate (Siddiqui et al., 2016). Because motility to and within the brain, and the phagocytosis-like mechanism used to consume brain tissue undoubtedly contribute to disease (Siddiqui et al., 2016), defining the molecular underpinnings of these specific cellular processes may reveal new and much-needed drug targets to combat this deadly pathogen.

## METHODS

### Cell and bacterial culture

*Naegleria gruberi* NEGM cells (a gift from Dr. Chan Fulton, Brandeis) were axenically cultured in M7 Media (See **table S1** for all reagents and recipes) in plug seal tissue culture-treated flasks (CELLTREAT, cat. no. 229330) and grown at 28°C. Cells were split into fresh media every 2-3 days. To induce differentiation (**Fig 1**), cells were centrifuged at 1500 RCF for 90 sec, pellets were resuspended in ice cold 2 mM Tris, and incubated in a 28°C water bath shaking at 100 RPM for approximately 90 min.

For phagocytosis assays, BL21 *E. coli* expressing superfolded-GFP (Pedelacq et al., 2006) were streaked from frozen glycerol stocks kept at -80 onto LB plates supplemented with 50 μg/ml Kanamycin. Broth cultures of LB + Kanamycin were inoculated from single colonies, and were grown overnight at 37°C shaking at 175 RPM in a New Brunswick Innova 4330 floor shaker. Cultures were then diluted 1:10 into fresh LB, and grown until the OD600 reached 0.4-0.7, at which point IPTG was added to a final concentration of 0.1 mM for 1 h.

### Transcript analysis and homolog identification

Microarray data from amoeba and flagellate populations were analyzed from experiments completed in (Fritz-Laylin and Cande, 2010). Each original biological replicate had been completed in duplicate, so the average for each biological replicate was first calculated. Then, the transcript abundance in flagellates (from the 80 min time point) was subtracted from the transcript abundance in amoebae (from the 0 min time point) for that biological replicate, and the average and standard deviation from the three replicates was calculated. Only the transcripts corresponding to actin or microtubule cytoskeletal genes were included in this analysis.

To expand upon the original *N. gruberi* cytoskeletal annotations ((Fritz-Laylin et al., 2010b), Table S5), database of additional proteins suspected to be present (e.g. additional Arp2/3 complex subunits, additional SCAR/WAVE complex subunits, WASH complex, etc.) was generated using protein sequences from human (obtained from pubmed) or *Dictyostelium discoideum* homologs (obtained from DictyBase: http://dictybase.org). The resulting database was used as a query to BLAST (Altschul et al., 1990) against the *N. gruberi* proteome, using default parameters except replacing the scoring matrix with the BLOSUM45 scoring matrix. We further validated protein identities through hmmscan searches using a gathering threshold and Pfam domains using the HMMER website (https://www.ebi.ac.uk/Tools/hmmer/search/hmmscan).

To assess the cytoskeletal repertoire of *Naegleria fowleri*, we compared an *N. fowleri* (ATCC 30863) protein database generated in Augustus (Herman et al., 2020) to *N. gruberi* actin and microtubule associated proteins (Fritz-Laylin et al., 2010b) using BLAST with default search parameters. The top *N. fowleri* hits identified by comparisons to *N. gruberi* were then compared using BLAST to the full *N. gruberi* protein library to establish best mutual BLAST hits. See **Table S2** for the full list of accession numbers.

### Multiple sequence alignments and inhibitor binding sites

Actin sequences from *Naegleria gruberi, N. fowleri*, and a diverse set of additional eukaryotes were collected (see **Table S3**) and aligned using T-Coffee (Notredame et al., 2000) with defaults (Blosum62 matrix, gap open penalty= -50, gap extension penalty= 0) in Jalview. Drug binding sites from the literature (Faulstich et al., 1993; Morton et al., 2000; Nair et al., 2008; Pospich et al., 2017) were mapped onto the rabbit muscle actin sequence, using only residues with evidence of direct interactions. It is important to note, however, that additional residues not highlighted in our MSA are likely required for the proper conformation of binding sites.

### Cell Morphology Assays and use of Inhibitors

For the experiments shown in Fig 3, flasks of cells were moved to room temperature on the experiment eve to prevent differentiation to flagellates during the experiment due to temperature fluctuation. 0.5 ml of cells were seeded (at ∼5×10^5^ cells/ml) into a 24 well plate preloaded with 0.5 ml fresh M7 media with inhibitors or controls, for final (1x) concentrations as follows: 5 μM LatB, 20 μM cytochalasin D,140 nM phalloidin, 2 μM jasplakinolide, 50 μM CK-666, 25 μM SMIFH2, 50 μM CK-689, 0.1% DMSO, and 0.09% ethanol. Concentrations were selected based on work in other systems (Velle and Campellone, 2018), and/or were tested at 10 fold dilutions within the range recommended in the company supplied data sheets in preliminary experiments. From preliminary experiments, the lowest concentration that had an observable phenotype in live cells was chosen, or in the case of cytochalasin D, jasplakinolide, and phalloidin, the highest concentrations we tested were chosen, which still did not produce any obvious phenotypes. Short time points (5-10 min) were selected for most experiments to avoid potential off-target effects of the inhibitors.

After 10 min of incubation with inhibitors or controls, cells were fixed and stained for microscopy following a protocol adapted from (Fritz-Laylin et al., 2010a). Media was aspirated from cells, 1 ml of microscopy fixative (see **Table S1** for all recipes and reagents) was added to each well, and 0.5 ml of cells were transferred to each of two coverslips (precoated with 0.1% polyethyleneimine) per condition. Cells were allowed to settle onto the coverslips while fixing. After 15 min of fixation, the fixative was aspirated from the wells, and replaced with 0.5 ml per well of permeabilization buffer. After 10 min of incubation, the permeabilization buffer was replaced with staining solution. After staining for 45 min, cells were washed 3 times in PEMBALG for ∼5 min each, and coverslips were mounted in Prolong Gold antifade reagent and allowed to cure overnight. The next morning, coverslips were sealed with nail polish, and labels were blinded prior to microscopic analysis.

Images for quantifying cell morphology were acquired using a Nikon Eclipse Ti2 microscope equipped with a pco.panda camera and a Plan Apo λ 100X Oil 1.45 NA objective. Alexa Fluor 488 labeled phalloidin and DAPI were imaged using a pE-300 MultiLaser LED light source (excitation: 460 nm or 400 nm; emission: 535 nm or 460 nm). Representative cells were imaged with Structured Illumination Microscopy (shown in Fig 2C and 3D) using a Nikon A1R-SIMe microscope equipped with an ORCA-Flash4.0 camera and an SR Apo TIRF 100x 1.49 NA objective.

Image analysis was performed on blinded datasets using Fiji (Schindelin et al., 2012) to view images as Z-stacks and to generate maximum intensity projections. Phenotypes were defined as follows: <u>Puncta</u>: A cell with at least 10 small (0.2-0.8 um diameter) F-actin rich foci in the cell counted as having puncta. <u>Ruffles</u>: Ruffles are actin-rich lamellar protrusions that resemble the ruffled edges of a lasagna noodle. <u>Filopodia-like protrusions</u>: Filopodia were defined as spaghetti-like protrusions from the main body of the cell that are longer than they are wide (less than 0.75 um thick, typically 0.2-0.5 um). <u>Area and Shape</u>: The maximum intensity projection was used to draw an outline of the arial footprint of the cell in ImageJ using the “kidney bean” tool. Measurements were taken to include the area and shape descriptors (of which, circularity was used). <u>Cortex measurements</u>: The cell cortex was measured by drawing a line (with a thickness of 3.25 μm) perpendicular to the cell edge using the line tool in imageJ. The center of the line was set to the cell edge so measurements could be normalized. Lines were preferentially drawn on non-curved sections of the cell, and a representative area of the membrane was chosen. The plot profile tool was used to generate pixel intensity plots, which were normalized to an area inside the cell that was set to 1.

### Cell Migration Assays

For the cell motility assays shown in Fig 4A-E, flasks of cells were moved to room temperature on the experiment eve to prevent differentiation to flagellates during the experiment. Cells were resuspended by slapping and swirling the flask, and 0.5 ml of cell suspension (∼10^5^ cells/ml) were added to each well of a 24 well plate (with a tissue culture treated surface). After allowing the cells to settle for 5 min, media was aspirated and cells were washed once with 0.5 ml Tris buffer (2 mM Tris, pH 7.2) per well. After the wash was aspirated, a final volume of 0.5 ml Tris buffer was added to the cells in each well. Cells were imaged using phase/contrast microscopy with a Nikon Eclipse Ti2 microscope equipped with a pco.panda camera and an Achromat 10x Ph1 ADL 0.25 NA objective, controlled by Nikon NIS Elements software. Cells were imaged using an acquisition rate of 5 sec/frame. After 5 min of imaging, an equal volume of 2x small molecule inhibitors or carriers alone (diluted in 2 mM Tris) were added to the wells of live amoebae. Imaging proceeded for an additional 5 min, and the exact time point inhibitors or controls were added was noted. File names were blinded, and 20 randomly selected cells from the center of the field of view at the time of treatment were chosen for tracking analysis. Tracked cells were only excluded if they left the field of view, or contacted another cell in a way that disrupted tracking (See SMIFH2 treatment, Movie S1). Cells were tracked using the mTrackJ plugin (Meijering et al., 2012) in Fiji, and the dataset was divided into “pre” and “post” treatment tracks for each cell with tracking data.

### Other Fluorescence Microscopy

Cells in the experiments in Fig 1B and S1 were spun down (1500 RCF/ 90 sec), resuspended in fixative (for 15 min), spun again, resuspended in permeabilization buffer (for 10 min), spun again, and resuspended in staining solution before transferring them to a 96-well glass-bottom plate. After 20 min, the staining solution was gently replaced with PBS. The cells in Fig 5A were fixed for flow cytometry, and imaged in a 96-well glass-bottom plate. The cells in Fig 5E were stained following the protocol used for cell morphology assays, except using AlexaFluor-568 phalloidin. The cells in Fig 4F-G were fixed, permeabilized, and stained in one step during imaging in a 35 mm tissue culture treated glass-bottom MatTek dish. Images in Fig 1B, S1, 4F-G, and 5 were taken using a Nikon Eclipse Ti2 microscope equipped with a pco.panda camera and a Plan Apo λ 100X Oil 1.45 NA objective or an Achromat 10x Ph1 ADL 0.25 NA objective (5A only).

### Electron Microscopy

NEGM cells were spun down at 1500 RCF for 90 sec, and the cell pellet was resuspended in Tris buffer. Cells were seeded onto aclar pieces (coated with 0.1% polyethyleneimine) in a 24 well plate, and allowed to settle before fixing in 2.5% glutaraldehyde + 100 mM cacodylate buffer (pH 7.2). After 20 min of fixation, cells were washed once, and dehydrated with a series of five minute ethanol washes at the following concentrations: 30%, 50%, 70%, 95%, 95%, 100%, 100%. Samples were then critical point dried, aclar pieces were mounted onto stubs using carbon tape, and sputter coated with 10 nm platinum. Samples were imaged on a Zeiss Supra 40VP scanning electron microscope.

### Flow Cytometry

#### Actin Polymer Content

Our protocol for measuring F-actin by flow cytometry (Kakley et al., 2018) was adapted for *Naegleria*. NEGM cells were treated with inhibitors for 15 minutes, spun down, and resuspended in flow cytometry fixative for 10 min. Cells were spun again, and pellets were resuspended in 50 ul PBS before transferring to flow cytometry tubes with permeabilization buffer and phalloidin. An extra ∼1 ml of PBS was added prior to flow cytometry. 20,000 cells were analyzed for each sample using a Fortessa flow cytometer controlled by FACS DIVA software. Analysis was completed using FlowJo.

#### Phagocytosis Assays

4 T75 tissue culture flasks of ∼60% confluent NEGM cells were transferred into Tris buffer 1 h prior to the experiment. Cells were then pooled and aliquoted into 125 ml erlenmeyer flasks (10 ml per flask) preloaded with 5 ml of Tris mixed with inhibitors or controls at 3x the final concentration (so 1x concentrations would be reached in the final volume of 15 ml). Then, *E. coli* induced to express sfGFP (protocol above) were spun down at 3000 RCF for 4 minutes and the pellet was resuspended in an equal volume of Tris to the starting culture. 750 ul of the bacterial suspension were added to each flask. Flasks were incubated while shaking at 28 C for 45 min. Cultures were then transferred to 15 ml conicals, spun at 1500 RCF/3 min, resuspended in 2 ml Tris buffer, spun again, and pellets were resuspended in 1 ml flow cytometry fixative and transferred to a 1.5 ml tube. After 15 min of fixing, cells were spun at 1500 RCF/2 min, and pellets were resuspended in 1 ml PBS and transferred to flow cytometry tubes. 30,000 cells were analyzed for each sample using a Fortessa flow cytometer controlled by FACS DIVA software. Analysis was completed using FlowJo.

### Data analysis and Statistics

The samples in Fig 4 D-H and Fig S5I were blinded before imaging, and were only unblinded after analyses on that experimental replicate were complete. Motility data were blinded after acquiring the data but before analysis. Flow cytometry samples were blinded before data acquisition, with the exception of phalloidin stained and unstained controls (Fig S5B), which were always run before the labor intensive analysis of the remaining controls and samples. All graphs were generated using Graphpad Prism software (version 8.2). Whenever applicable, SuperPlots (Lord et al., 2020) were employed to show data on each individual cell (smaller, gray symbols), while also displaying averages from each experimental replicate (larger, colorful symbols). The experimental replicates were used to determine the mean, standard deviation, and statistical significance (with an ordinary ANOVA and Tukey’s multiple comparison test).

## ACKNOWLEDGEMENTS

We thank Chandler Fulton (Brandeis University) for NEGM cells and Katherine Dorfman (University of Massachusetts) for sfGFP-expressing *E. coli*. We thank James Chambers for assistance with Structured Illumination Microscopy, which was performed at the Light Microscopy Facility and Nikon Center of Excellence at the Institute for Applied Life Sciences, UMass Amherst with support from the Massachusetts Life Sciences Center. We thank Amy S Burnside, the Director of the Flow Cytometry Core Facility at UMass Amherst/IALS, for technical expertise, and Shane Hussey for assistance with flow cytometry. We also thank Kasia Hammar and the Marine Biological Laboratory central microscopy center staff for assistance with Electron Microscopy. We thank Simon Shulman for collecting preliminary data for this project. We also thank Samuel Lord (University of California San Francisco), Patricia Wadsworth (UMass), Kenneth Campellone (University of Connecticut), and Edgar Medina (UMass) for providing helpful feedback on this manuscript. This work was supported by The National Institutes of Health (grant 1R21AI139363 from the National Institute of Allergy and Infectious Diseases), The Pew Charitable Trust, and an Excellence in Biomedical Research Award from the Smith Family Foundation.

## SUPPLEMENTAL MOVIES

**Movie S1: NEGM motility is altered by different small molecule inhibitors.** Crawling NEGM amoebae were imaged using phase/contrast microscopy at a rate of 5s/frame. Cells were treated with the indicated inhibitors or controls at the 5 min mark, and imaging proceeded for an additional 5 min.

**Movie S2: CK-666 induces actin-rich filopodial protrusions.** Cells were imaged using DIC microscopy at a frame rate of 1s/frame, and treated with CK-666 during imaging. Cells were simultaneously fixed, permeabilized, and stained, with DAPI to detect DNA (magenta), and phalloidin to detect F-actin (green).

## SUPPLEMENTAL TABLES

**Table S1:** Reagents and recipes used in this study.

|  | Reagent | Company | Cat No | Working concentration |
| --- | --- | --- | --- | --- |
| Small molecule inhibitors | Latrunculin B | Millipore | 4280201MG | 5 μM |
| | Jasplakinolide | Molecular Probes/Invitrogen | J7473 | 2 μM |
|  | Phalloidin | Invitrogen | P3457 | 140 nM |
| | Cytochalasin D | Gibco/Life Technologies | PHZ1063 | 20 μM |
| | CK-666 | Calbiochem/Sigma | 182515 | 50 μM |
| | CK-689 | Calbiochem/Sigma | 182517 | 50 μM |
| | SMIFH2 | Tocris Bioscience | 440110 | 25 μM |
| Stains/cell dyes | 4',6-diamidino-2-phenylindole (DAPI) | Life Technologies | D1306 | 1 μg/ml |
|  | Alexa Fluor™ 488 Phalloidin | Invitrogen | A12379 | 6.6 nM - 66 nM |
|  | Alexa Fluor™ 568 Phalloidin | Invitrogen | A12380 | 6.6 nM - 66 nM |
|  | Tubulin Tracker Deep Red | Life Technologies | T34077 | 1X (see user guide) |
|  | ProLong Gold Antifade reagent + DAPI | Invitrogen | P36935 | N/A |
| Other Chemicals | Reagent | Company | Cat No |  |
|  | Agar | APEX | 20-248 |  |
|  | Bovine serum albumin (BSA) | Fisher | BP1600-1 |  |
|  | Dimethyl Sulfoxide (DMSO) | Sigma | D2650-100 ml |  |
|  | EGTA | Sigma | E4378-100g |  |
|  | Fetal bovine serum (FBS) | Atlanta Biologicals | S11150 (Lot Number K18142) |  |
|  | Gelatin from cold water fish skin | Sigma | G7765-200ml |  |
|  | Glucose | Sigma | G5767-5kg |  |
|  | Glutaraldehyde (25%) | Electron Microscopy Sciences | 16220 |  |
|  | Kanamycin sulfate | Fisher | BP906-5 |  |
|  | Lysogeny broth, Miller | Sigma | L3152-1kg |  |
|  | L-methionine | Sigma | M5308-25g |  |
|  | Lysine | Sigma | L5626-1kg |  |
|  | Magnesium sulfate heptahydrate (MgSO4 7H2O) | Sigma | 230391-500g |  |
|  | NP-40 Alternative | Millipore | 492016-500ml |  |
|  | Paraformaldehyde (16%) | Electron Microscopy Sciences | 15710 |  |
|  | Penicillin Streptomycin Solution | Sigma | P4333-100ML |  |
|  | PIPES | Sigma | p6757-500g |  |
|  | Polyethyleneimine | Sigma | p3143-100 ml |  |
|  | Potassium chloride (KCl) | Fisher | p217-500 |  |
|  | Potassium phosphate monobasic (KH2PO4) | Sigma | p0662-1kg |  |
|  | Sodium Azide (N3Na) | Sigma | S2002-100g |  |
|  | Sodium Cacodylate Trihydrate | Sigma | C4945-100g |  |
|  | Sodium chloride (NaCl) | Fisher | BP358-1 |  |
|  | Sodium Phosphate dibasic (Na2HPO4) | Sigma | s9763-1kg |  |
|  | Sodium Phosphate Monobasic | Sigma | s9638-500g |  |
|  | Sodium phosphate monobasic monohydrate (NaH2PO4 H2O) | Sigma | s9638-500g |  |
|  | Sucrose | Fisher | s5-500 |  |
|  | Tris | Fisher | BP152-5 |  |
|  | Yeast extract | Fisher | BP1422-2 |  |
| Recipes | Solution | Recipe |  |  |
| | LB Broth | 25 g/L LB broth, +/- 50 μg/ml Kanamycin | | |
| | LB Plates | 25 g/L LB broth + 20 g/L Agar +/- 50 μg/ml Kanamycin | | |
|  | M7 Media | 10% FBS + 1x Penicillin Streptomycin Solution + 45 mg/L L-methionine + 5 g/L yeast extract + 5.4 g/L glucose + 2% M7 Buffer |  |  |
|  | M7 Buffer | 18.1 g/L KH2PO4 + 25 g/L Na2HPO4 |  |  |
|  | PBS | 0.137 M NaCl + 2.7 mM KCl + 10 mM Na2HPO4 + 1.8 mM KH2PO4 |  |  |
|  | 500 mM Sodium Phosphate Buffer, pH 7.2 | 280 ml 0.5 M NaH2PO4 + 720 ml 0.5 M Na2HPO4 |  |  |
|  | PEM | 100 mM PIPES + 1 mM EGTA + 0.1 mM MgSO4 |  |  |
|  | PEMBALG | 100 mM PIPES + 1 mM EGTA + 0.1 mM MgSO4 + 0.1% BSA + 0.1% Sodium Azide + 100 mM Lysine + 0.5% Fish Gelatin |  |  |
|  | Electron Microscopy Fixation Buffer | 2.5% Glutaraldehyde + 100 mM Sodium Cacodylate |  |  |
|  | Flow Cytometry Fixation Buffer | 4% PFA + 25 mM Sodium Phosphate Buffer + 65 mM Sucrose |  |  |
|  | Microscopy Fixation Buffer | 1.8% PFA + 65 mM sucrose + 25 mM sodium phosphate buffer |  |  |
|  | Flow Cytometry Permeabilization Buffer | 0.1% NP-40 Alternative in PBS |  |  |
|  | Microscopy Permeabilization Buffer | 0.1% NP-40 Alternative + 6.6 nM Alexa Fluor 488 (or 568, Fig 5 only) Phalloidin in PEM |  |  |
|  | Flow Cytometry Stain (Fig 3B-C, S5D-H) | 66 nM Alexa Fluor 488 Phalloidin in PBS |  |  |
| | Microscopy Stain (Fig 1B, S1) | 66 nM Alexa Fluor 488 Phalloidin + 1x Tubulin Tracker Deep Red + 1 μg/ml DAPI in PEMBALG | | |
| | Microscopy Stain (Fig 2C, 3D, S5A, S6) | 66 nM Alexa Fluor 488 Phalloidin + 1 μg/ml DAPI in PEMBALG | | |
| | Microscopy Stain (Fig 5E) | 66 nM Alexa Fluor 568 Phalloidin + 1 μg/ml DAPI in PEMBALG | | |
| | 2x Microscopy Fix, Permeabilization Buffer, and Stain (Fig 4 F-G) | 1.8% PFA + 65 mM sucrose + 25 mM sodium phosphate buffer + 0.1% NP-40 Alternative + 132 nM Alexa Fluor 488 Phalloidin + 1 μg/ml | | |

**Table S2:**
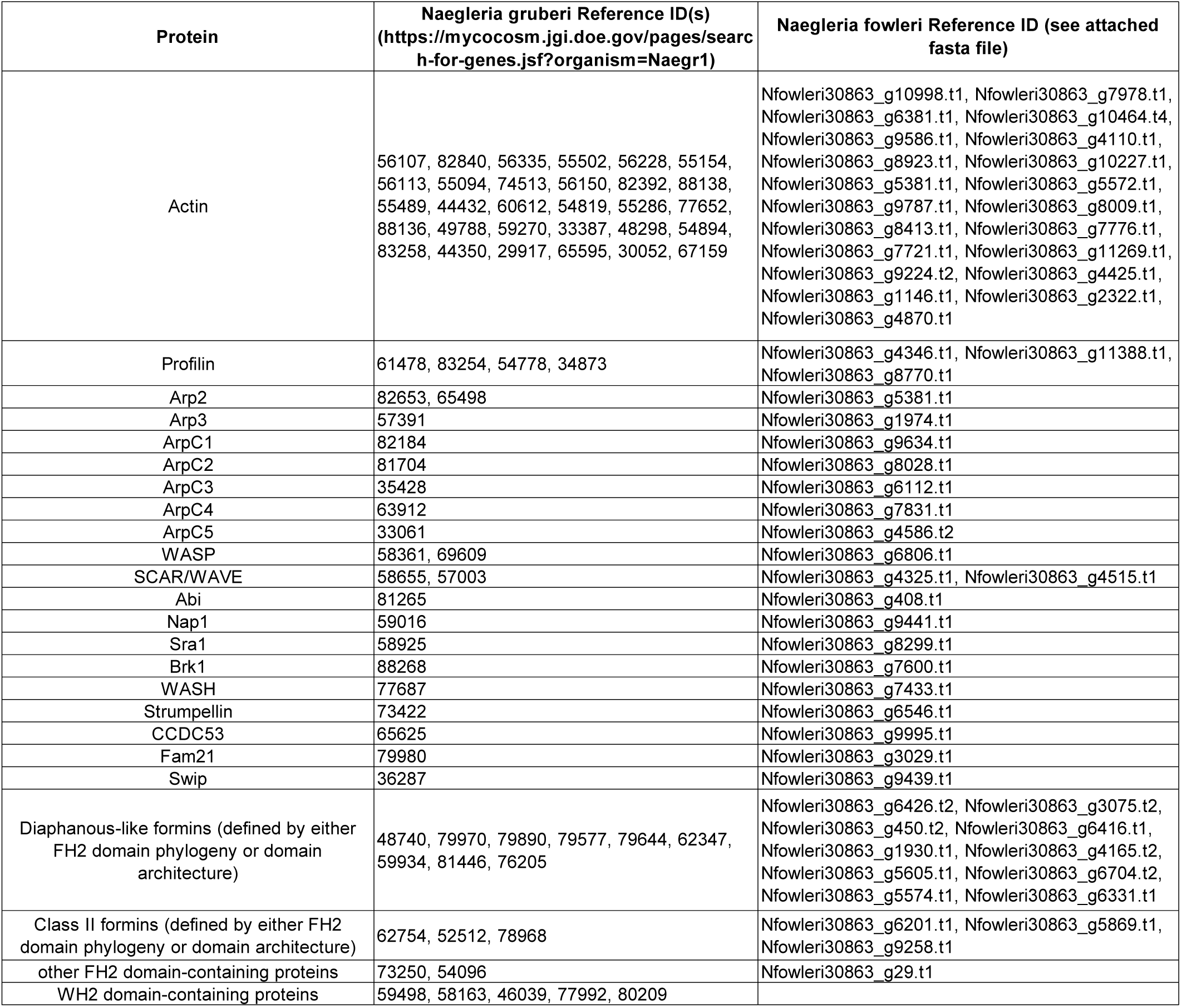
*Naegleria* cytoskeletal genes.

| Protein | Naegleria gruberi Reference ID(s)<br>(https://mycocosm.jgi.doe.gov/pages/search-for-genes.jsf?organism=Naegr1) | Naegleria fowleri Reference ID (see attached<br>fasta file) |
| --- | --- | --- |
| Actin | 56107, 82840, 56335, 55502, 56228, 55154,<br>56113, 55094, 74513, 56150, 82392, 88138,<br>55489, 44432, 60612, 54819, 55286, 77652,<br>88136, 49788, 59270, 33387, 48298, 54894,<br>83258, 44350, 29917, 65595, 30052, 67159 | Nfowleri30863_g10998.t1, Nfowleri30863_g7978.t1,<br>Nfowleri30863_g6381.t1, Nfowleri30863_g10464.t4,<br>Nfowleri30863_g9586.t1, Nfowleri30863_g4110.t1,<br>Nfowleri30863_g8923.t1, Nfowleri30863_g10227.t1,<br>Nfowleri30863_g5381.t1, Nfowleri30863_g5572.t1,<br>Nfowleri30863_g9787.t1, Nfowleri30863_g8009.t1,<br>Nfowleri30863_g8413.t1, Nfowleri30863_g7776.t1,<br>Nfowleri30863_g7721.t1, Nfowleri30863_g11269.t1,<br>Nfowleri30863_g9224.t2, Nfowleri30863_g4425.t1,<br>Nfowleri30863_g1146.t1, Nfowleri30863_g2322.t1,<br>Nfowleri30863_g4870.t1 |
| Profilin | 61478, 83254, 54778, 34873 | Nfowleri30863_g4346.t1, Nfowleri30863_g11388.t1,<br>Nfowleri30863_g8770.t1 |
| Arp2 | 82653, 65498 | Nfowleri30863_g5381.t1 |
| Arp3 | 57391 | Nfowleri30863_g1974.t1 |
| ArpC1 | 82184 | Nfowleri30863_g9634.t1 |
| ArpC2 | 81704 | Nfowleri30863_g8028.t1 |
| ArpC3 | 35428 | Nfowleri30863_g6112.t1 |
| ArpC4 | 63912 | Nfowleri30863_g7831.t1 |
| ArpC5 | 33061 | Nfowleri30863_g4586.t2 |
| WASP | 58361, 69609 | Nfowleri30863_g6806.t1 |
| SCAR/WAVE | 58655, 57003 | Nfowleri30863_g4325.t1, Nfowleri30863_g4515.t1 |
| Abi | 81265 | Nfowleri30863_g408.t1 |
| Nap1 | 59016 | Nfowleri30863_g9441.t1 |
| Sra1 | 58925 | Nfowleri30863_g8299.t1 |
| Brk1 | 88268 | Nfowleri30863_g7600.t1 |
| WASH | 77687 | Nfowleri30863_g7433.t1 |
| Strumpellin | 73422 | Nfowleri30863_g6546.t1 |
| CCDC53 | 65625 | Nfowleri30863_g9995.t1 |
| Fam21 | 79980 | Nfowleri30863_g3029.t1 |
| Swip | 36287 | Nfowleri30863_g9439.t1 |
| Diaphanous-like formins (defined by either<br>FH2 domain phylogeny or domain<br>architecture) | 48740, 79970, 79890, 79577, 79644, 62347,<br>59934, 81446, 76205 | Nfowleri30863_g6426.t2, Nfowleri30863_g3075.t2,<br>Nfowleri30863_g450.t2, Nfowleri30863_g6416.t1,<br>Nfowleri30863_g1930.t1, Nfowleri30863_g4165.t2,<br>Nfowleri30863_g5605.t1, Nfowleri30863_g6704.t2,<br>Nfowleri30863_g5574.t1, Nfowleri30863_g6331.t1 |
| Class II formins (defined by either FH2<br>domain phylogeny or domain architecture) | 62754, 52512, 78968 | Nfowleri30863_g6201.t1, Nfowleri30863_g5869.t1,<br>Nfowleri30863_g9258.t1 |
| other FH2 domain-containing proteins | 73250, 54096 | Nfowleri30863_g29.t1 |
| WH2 domain-containing proteins | 59498, 58163, 46039, 77992, 80209 |  |

**Table S3:** Eukaryotic actin sequences.

| Organism | Database and Reference ID |
| --- | --- |
| *Naegleria gruberi* | JGI: 56107 |
| *Naegleria fowleri* | Attached FASTA file:<br>Nfowleri30863_g10998.t1 |
| *Trypanosoma cruzi* (strain CL Brener) | NCBI Reference Sequence:<br>XP_807262.1 |
| *Giardia intestinalis* | GenBank: ESU35627.1 |
| *Trichomonas vaginalis* | NCBI Reference Sequence:<br>XP_001305234.1 |
| *Chlamydomonas reinhardtii* (NAP1) | NCBI Reference Sequence:<br>XP_001703266.1 |
| *C. reinhardtii* (IDA5) | NCBI Reference Sequence:<br>XP_001699068.1 |
| *Ectocarpus siliculosus* | GenBank: CBJ30601.1 |
| *Toxoplasma gondii* | NCBI Reference Sequence:<br>XP_002369663.1 |
| *Plasmodium falciparum* (PfActin-2) | UniProtKB/Swiss-Prot:<br>P14883.1 |
| *P. falciparum* (PfActin) | GenBank: KOB88525.1 |
| *Thalassiosira pseudonana* | NCBI Reference Sequence:<br>XP_002294917.1 |
| *Tetrahymena thermophila* | UniProtKB/Swiss-Prot:<br>P10992.3 |
| *Dictyostelium discoideum* | UniProtKB/Swiss-Prot:<br>P07830.2 |
| *Entamoeba histolytica* | NCBI Reference Sequence:<br>XP_001913457.1 |
| *Acanthamoeba castellanii* | NCBI Reference Sequence:<br>XP_004337269.1 |
| *Schizosaccharomyces pombe* | GenBank: BAA12315.1 |
| *Saccharomyces cerevisiae* | NCBI Reference Sequence:<br>NP_116614.1 |
| *Oryctolagus cuniculus* (muscle) | UniProtKB/Swiss-Prot:<br>P68135.1 |
| *O. cuniculus* Arp1 | NCBI Reference Sequence:<br>XP_002718680.1 |

## SUPPLEMENTAL FILES

**File S1: *N. fowleri* protein sequences**

**File S2: Eukaryotic actin sequences**

---

**Fig S1.**
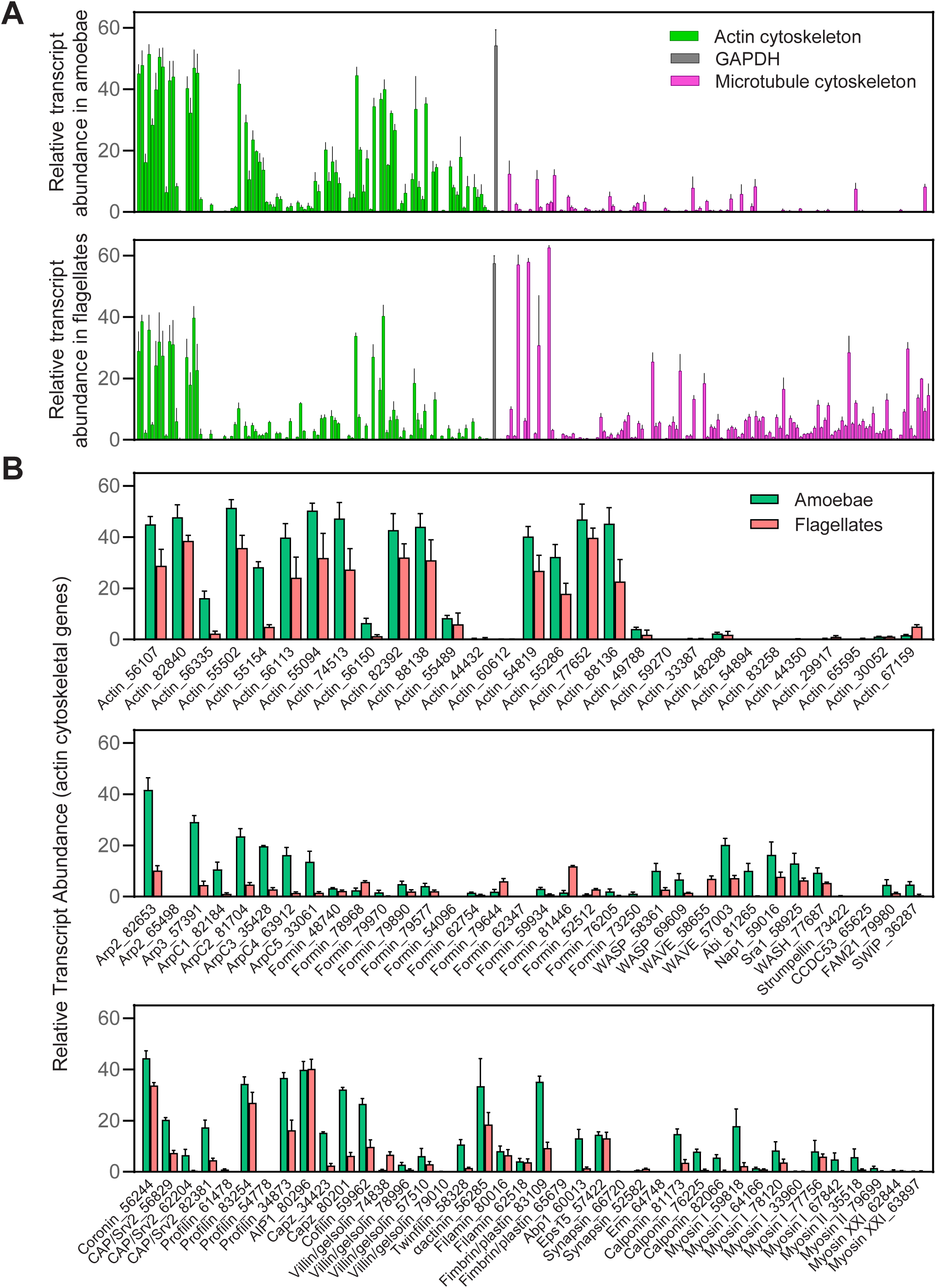
Actin and microtubule cytoskeletal gene expression differs between *Naegleria* amoebae and flagellates. (**A**) The relative transcript abundance for actin cytoskeletal genes (green), microtubule cytoskeletal genes (magenta), and GAPDH (gray) were calculated using expression data collected from amoeba (0 min into differentiation) or flagellate (80 min into differentiation) populations (original data from Fritz-Laylin and Cande, 2010). The relative transcript abundance in flagellates was subtracted from the level in amoebae to generate the graphs shown in Fig 2B-C. (**B**) The actin cytoskeletal transcript levels shown in panel A were organized such that the relative abundance in amoebae (jade green) and flagellates (salmon) were side-by-side for each of the indicated genes (numbers are in reference to the JGI accession numbers). Actins are shown in the top panel, nucleators and NPFs are grouped in the middle panel, and other actin binding proteins and motors are shown in the bottom panel. For all graphs, each bar represents the average relative transcript abundance from 3 independent experiments +/- SD.

**Fig S2.**
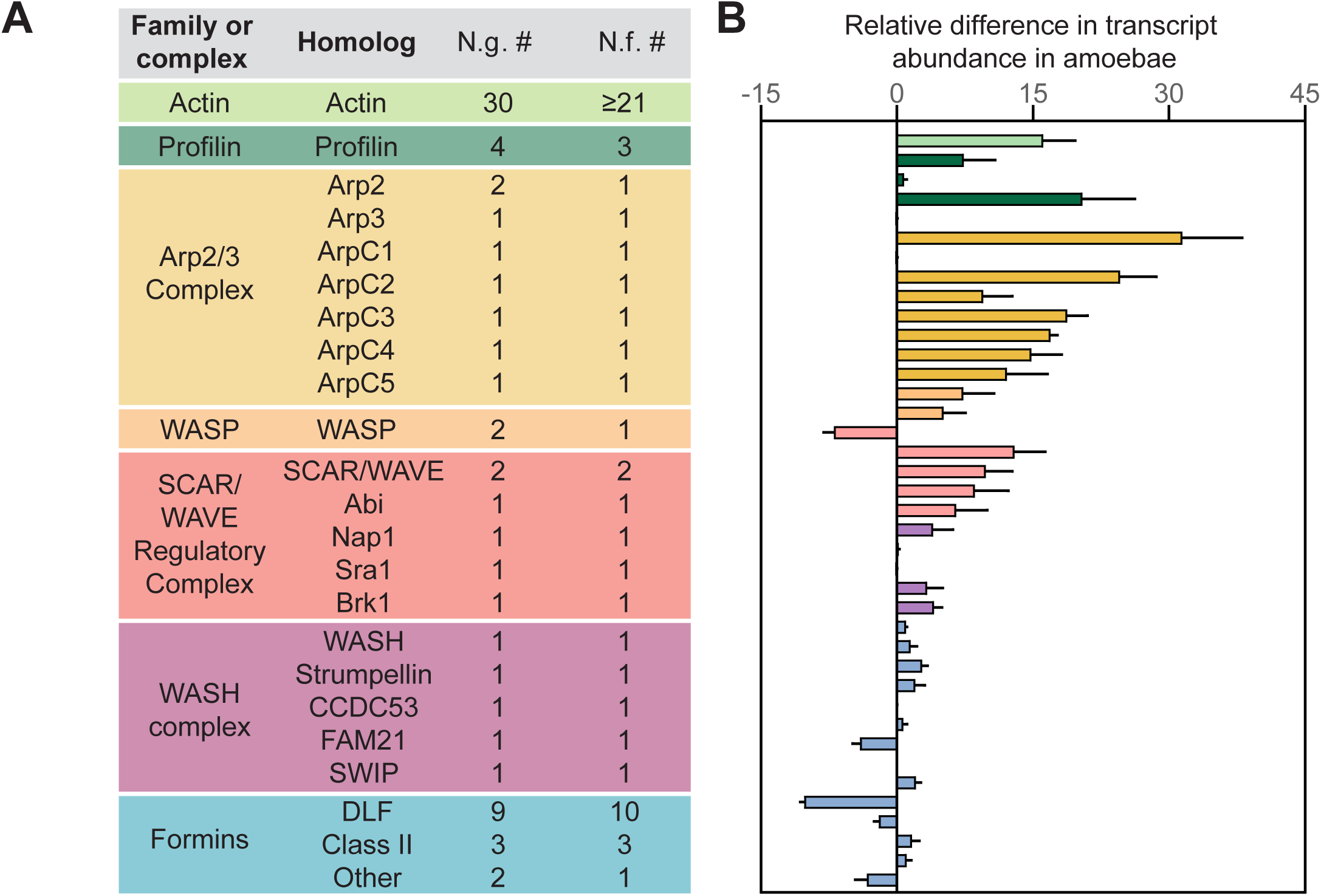
*Naegleria* species encode an extensive repertoire of actin cytoskeletal regulators. (**A**) *Naegleria gruberi (N.g.)* and *Naegleria fowleri (N.f.)* genomes each encode a similar cellular complement of actin cytoskeletal proteins. The first column shows the name of the protein family or complex, the second includes the name of the mammalian homolog (with the exception of Class II formins, which are often found in plants, and Diaphanous Like Formins (DLF), which encompass all diaphanous related formins), and the final columns indicate the number of genes found encoding these proteins in *N. gruberi* and *N. fowleri*, respectively. See Table S2 for accession numbers and sequences. We note that *N. gruberi* also possesses 5 WH2-domain containing proteins not listed above, and no homolog of WASP Interacting Protein was identified in either *Naegleria* genome. (**B**) The same graph from Figure 2C is shown for side-by-side comparisons of expression data with the proteins listed in panel A. Each bar represents the average relative change in transcript abundance between amoebae and flagellates from 3 experiments +/- Standard Deviation (SD).

**Fig S3.**
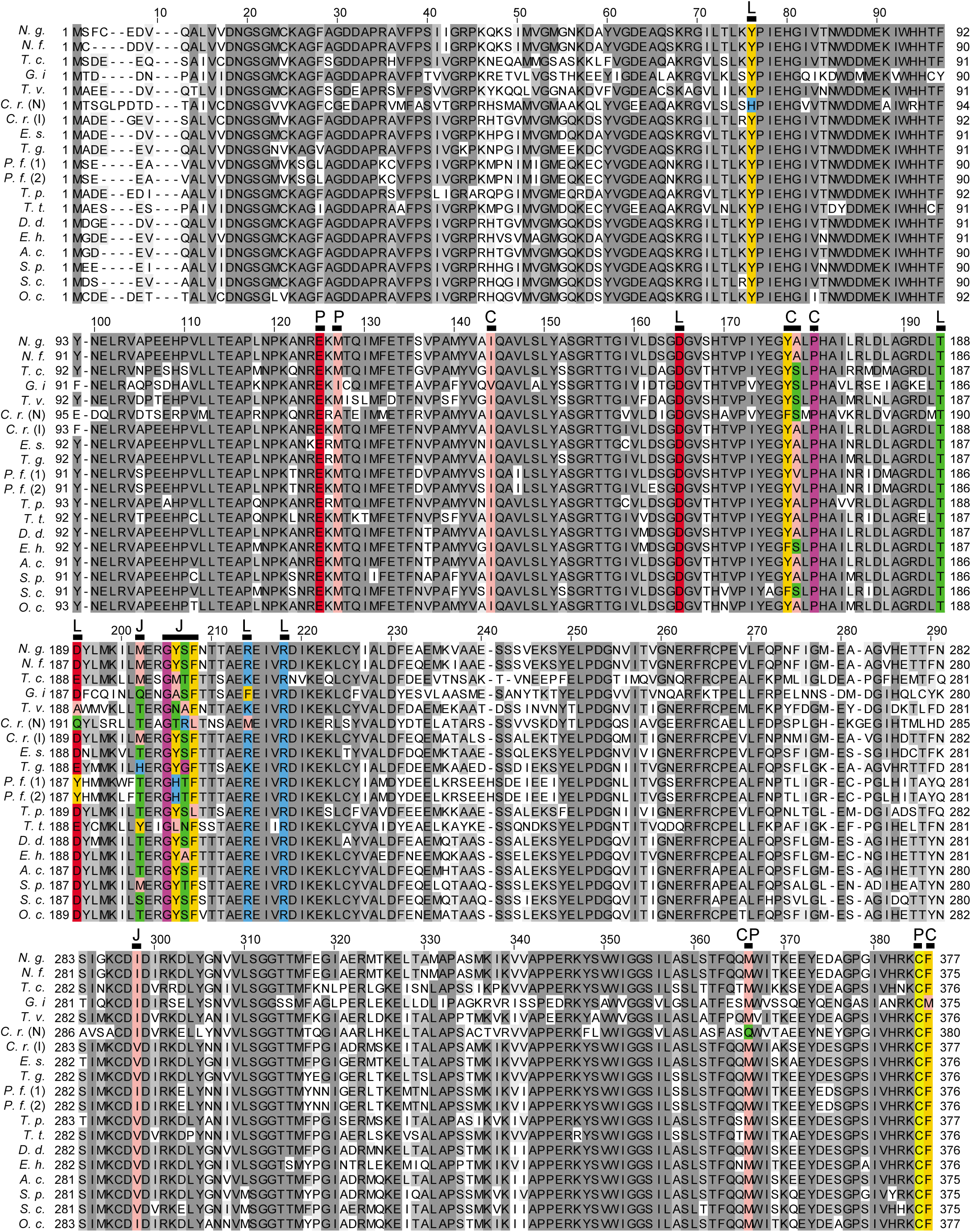
*Naegleria* actin retains conserved binding sites for small molecule inhibitors of actin dynamics. A multiple sequence alignment was generated to compare *Naegleria gruberi’*s (N.g) most highly expressed actin protein sequence with other eukaryotic actins (see Table S3 for sequences and ID numbers). This alignment includes actin sequences from: other Discobids, *N. fowleri* (N.f.) and *Trypanasoma cruzi* (T.c.); Metamonads, *Giardia lamblia* (G.i.) and *Trichomonas vaginalis* (T.v.); Plants, *Chlamydomonas reinhardtii* (C.r., including NAP1 (N) and IDA5 (I)) and *Ectocarpus siliculosus* (E.s.); Stramenopiles, *Thalassiosira pseudonana* (T.p.); Alveolates, *Toxoplasma gondii* (T.g.), *Plasmodium falciparum* (P.f, including actin-1 and actin-2), and *Tetrahymena thermophila* (T.t.); Amoebozoa, *Dictyostelium discoideum* (D.d.), *Entamoeba histolytica* (E.h), and *Acanthamoeba castellanii* (A.c.); and Opisthokonts, *Saccharomyces cerevisiae* (S.c.), *Schizosaccharomyces pombe* (S.p.), and *Oryctolagus cuniculus* (O.c.). Sequences were aligned using T-Coffee (Notredame et al., 2000) with defaults (Blosum62 matrix, gap open penalty= -50, gap extension penalty= 0) in Jalview, and residues were colored based on conservation (grayscale), or to highlight differences in amino acids within drug binding sites (ILVAM: salmon; FWY: tangerine; KRH: blue; DE: red; STNQ: lime; PG: raspberry; C: yellow). Bars indicate important residues for drug binding, and binding sites are abbreviated as follows: L, Latrunculin; P, Phalloidin; C, Cytochalasin; J, Jasplakinolide.

**Fig S4.**
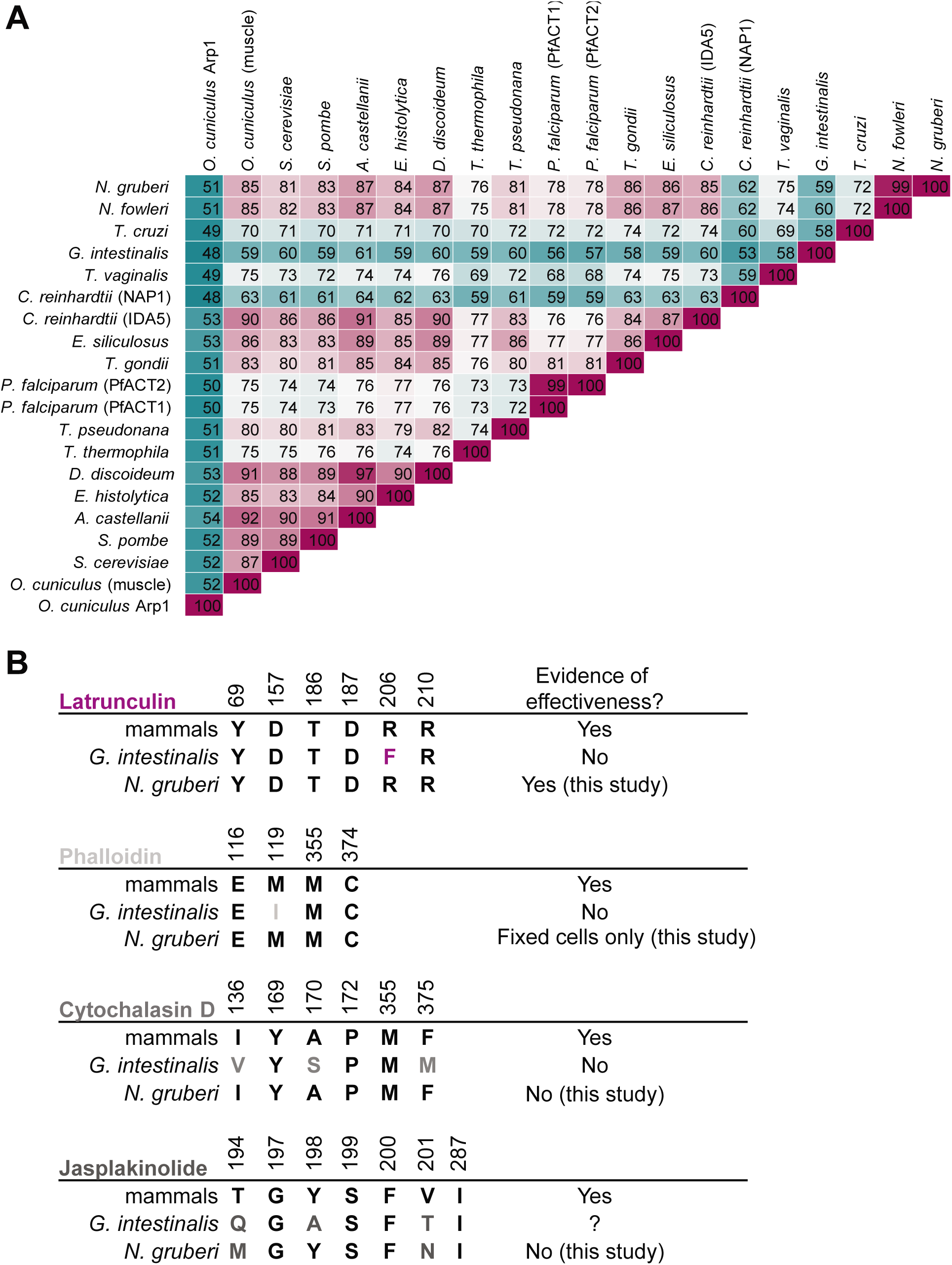
Binding sites for LatB, Phalloidin, and CytoD are identical between mammalian and *Naegleria* actins. (**A**) The alignment shown in Fig S3 was used to calculate the percent identity of actin protein sequences from representative eukaryotes. Values indicating higher conservation are highlighted in purple and values indicating less conservation are highlighted in teal. Arp1 from rabbit was included for comparison to an actin-related protein. (**B**) Drug binding sites on mammalian actin (Morton et al., 2000; Faulstich et al., 1993; Nair et al., 2008; Pospich et al., 2017) are shown in comparison to *Giardia* actin and *Naegleria* actin. Conserved residues are shown in black, and the numbers above each column indicate the location of the residue in the mammalian sequence. *Giardia* actin is shown as an example, due to its low percent identity to mammalian actin, as well as the documented ineffectiveness of actin inhibitors (Paredez et al., 2011). Question marks indicate unknown information.

**Fig S5.**
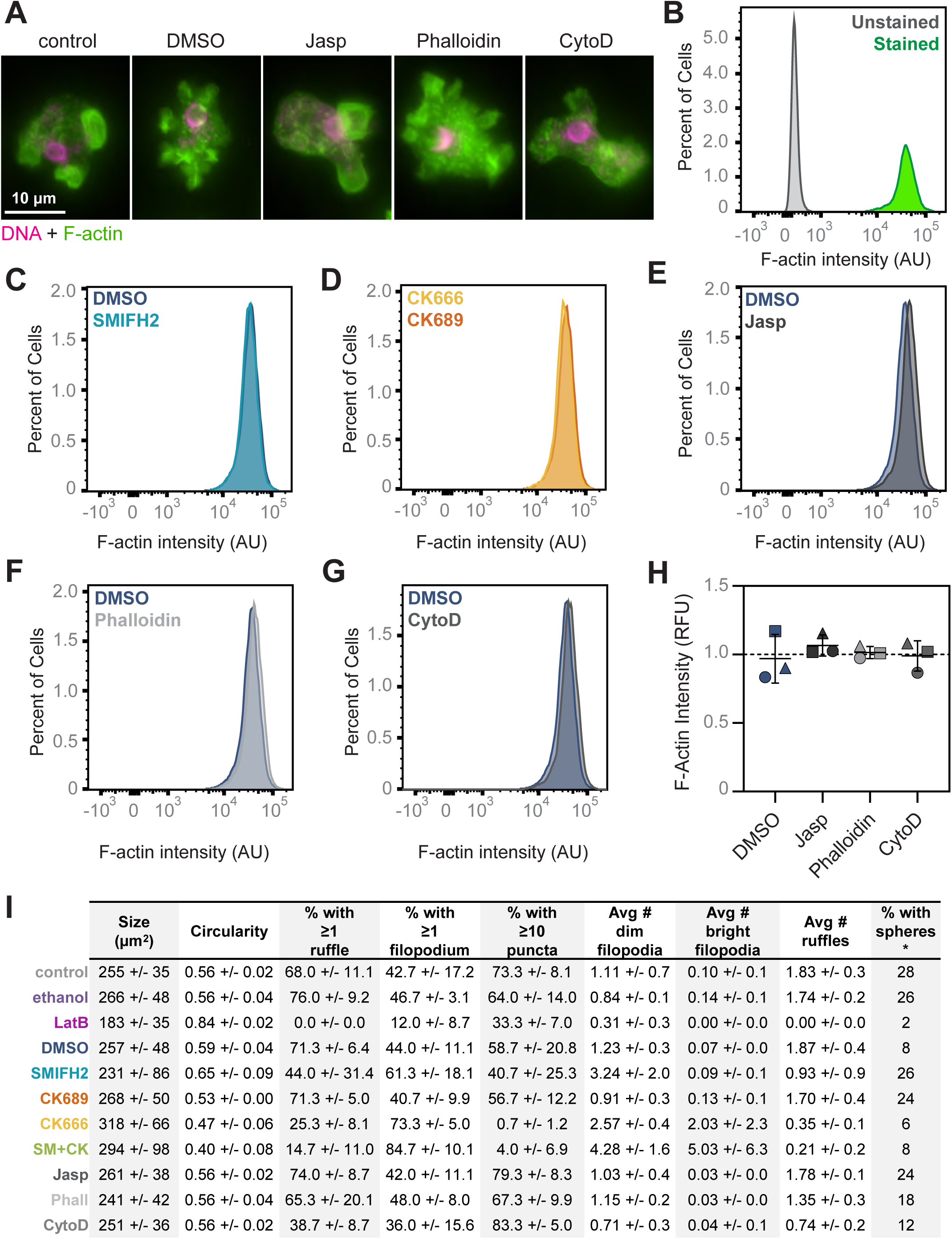
Neither cell morphology nor actin polymer content is affected by Jasplakinolide, Phalloidin, or Cytochalasin. (**A**) NEGM cells were incubated in media +/- inhibitors or controls for 10 min, then fixed and stained with Alexa-488 labeled phalloidin to detect F-actin (green) and DAPI to label DNA (magenta) prior to imaging using widefield fluorescence microscopy. Representative cells are shown. (**B-G**) NEGM cells treated as in A were stained only with Alexa-488 labeled phalloidin (with the exception of an unstained control, shown in B) prior to analysis by flow cytometry. Representative histograms of F-actin staining intensity compare drug treatments to respective controls for 1 of 3 biological replicates. (**H**) Average intensities calculated from E-G were normalized to the stained control (B). Each point represents the average normalized fluorescence intensity of F-actin staining, and lines indicate the mean of three experimental replicates +/- SD. Statistical significance was determined using an ordinary ANOVA followed by Tukey’s multiple comparison test. (**I**) Images like those shown in A were analyzed with the data set presented in Figure 4. Values are averages +/-SD from 3 independent replicates, each of which encompassed 50 cells per treatment condition. No statistical differences were found comparing Jasplakinolide, Phalloidin, or Cytochalasin D treatments to controls using an ordinary ANOVA followed by Tukey’s multiple comparison test. *The percent of cells with actin spheres was calculated for one representative dataset, and due to large variability between control samples, was not quantified for the remaining data sets.

**Fig S6.**
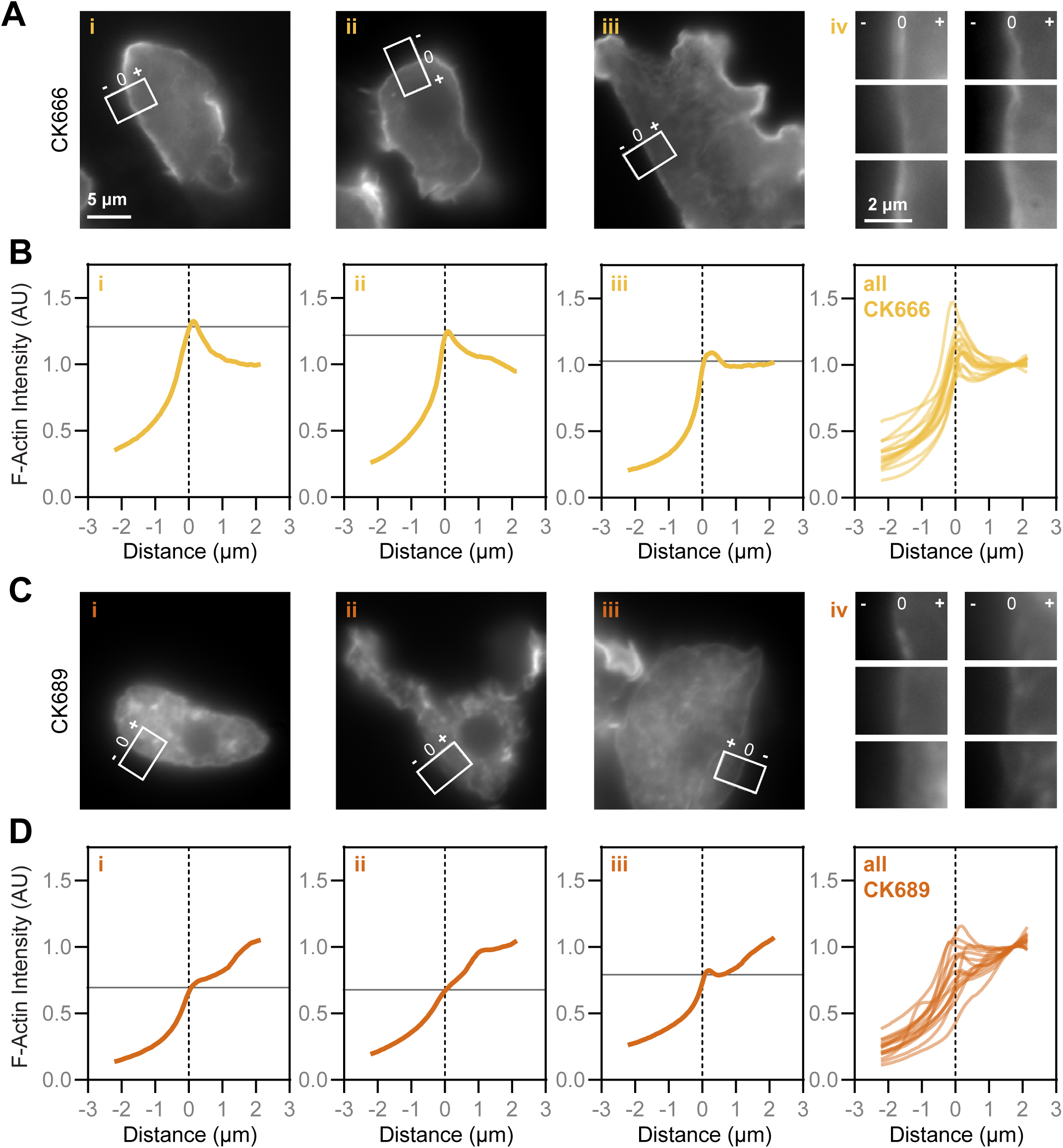
CK-666 treatment enhances the relative concentration of cortical to intracellular F-actin. (**A**) 15 NEGM cells treated with the Arp2/3 complex inhibitor CK-666 from the experiments described in Figure 4 were imaged using widefield fluorescence microscopy. A line with a width of 50 px, representing 3.25 μm was drawn perpendicular to the cell edge on a random, relatively straight part of each cell, in a z plane in which the cell edge was in focus. Panels i-iii show examples of these regions on cells, with one example from each of the three biological replicates analyzed. Panel iv shows the regions of interest of six additional examples. (**B**) The pixel intensities along the long axis of each box shown in A were normalized to the average pixel intensity of an area inside the cell, which was set to 1. These normalized intensities were plotted against the distance, with 0 indicating the cell edge (dashed line). The horizontal line indicates the maximum fluorescence intensity at the cell edge. The first 3 panels (i-iii) correspond to the images in A i-iii, and the rightmost “all” panel depicts all the data used to generate the plot in Fig 3J. (**C-D**) C and D mirror A and B, respectively, using the inactive control CK-689.

**Fig S7.**
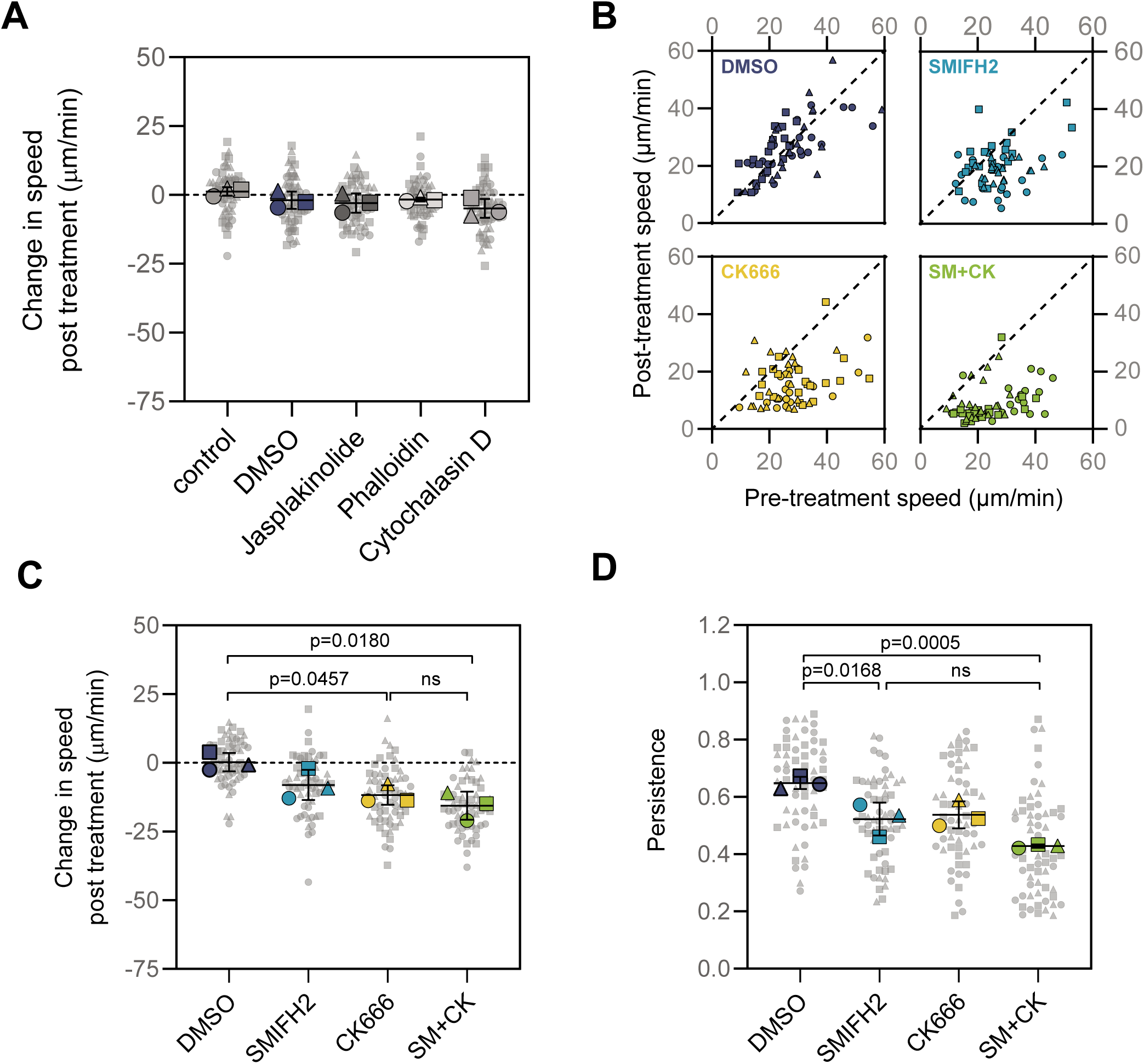
Jasplakinolide, Phalloidin, and Cytochalasin D do not impair cell speed, and combining Arp2/3 and formin inhibitors does not further impair cell motility. (**A**) In addition to the inhibitors shown in Fig 5, each experimental replicate also included cells treated with Jasplakinolide, Phalloidin, and CytoD. Each small, gray symbol represents the change in speed of a single cell after treatment, while larger symbols represent the averages from each experimental replicate, coordinated by shape (circles, squares, and triangles). (**B**) Cells were imaged as in Fig 5; after 5 minutes of imaging in buffer, cells were treated with DMSO, SMIFH2, CK-666, or a cocktail of SMIFH2 and CK-666 (SM+CK), and imaged for 5 additional min. 20 randomly selected cells from the center of the field of view at the time of treatment were tracked to calculate average speeds before and after treatment. Each point represents the speed of a cell before treatment (plotted on the x axis) and after treatment (plotted on the y axis). 3 experimental replicates are indicated using different shapes (squares, triangles, and circles). The dashed line has a slope of 1, indicating where cells would fall if the speed was unchanged. (**C**) The data collected from experiments in B were used to calculate the change in cell speed post-treatment. Each smaller gray symbol represents a single cell, while larger symbols represent the averages from each experimental replicate, with shapes representing independent replicates. (**D**) Directional persistence was calculated for each post-treatment cell tracked in B by dividing the maximum displacement from the start of the track by the total path length. Each small gray symbol represents the persistence of a single cell, while larger symbols represent experimental averages. For A, C, and D, Black lines indicate the mean +/- SD calculated from the 3 experimental replicates, and statistical significance was determined using an ordinary ANOVA followed by Tukey’s multiple comparison test.

**Fig S8.**
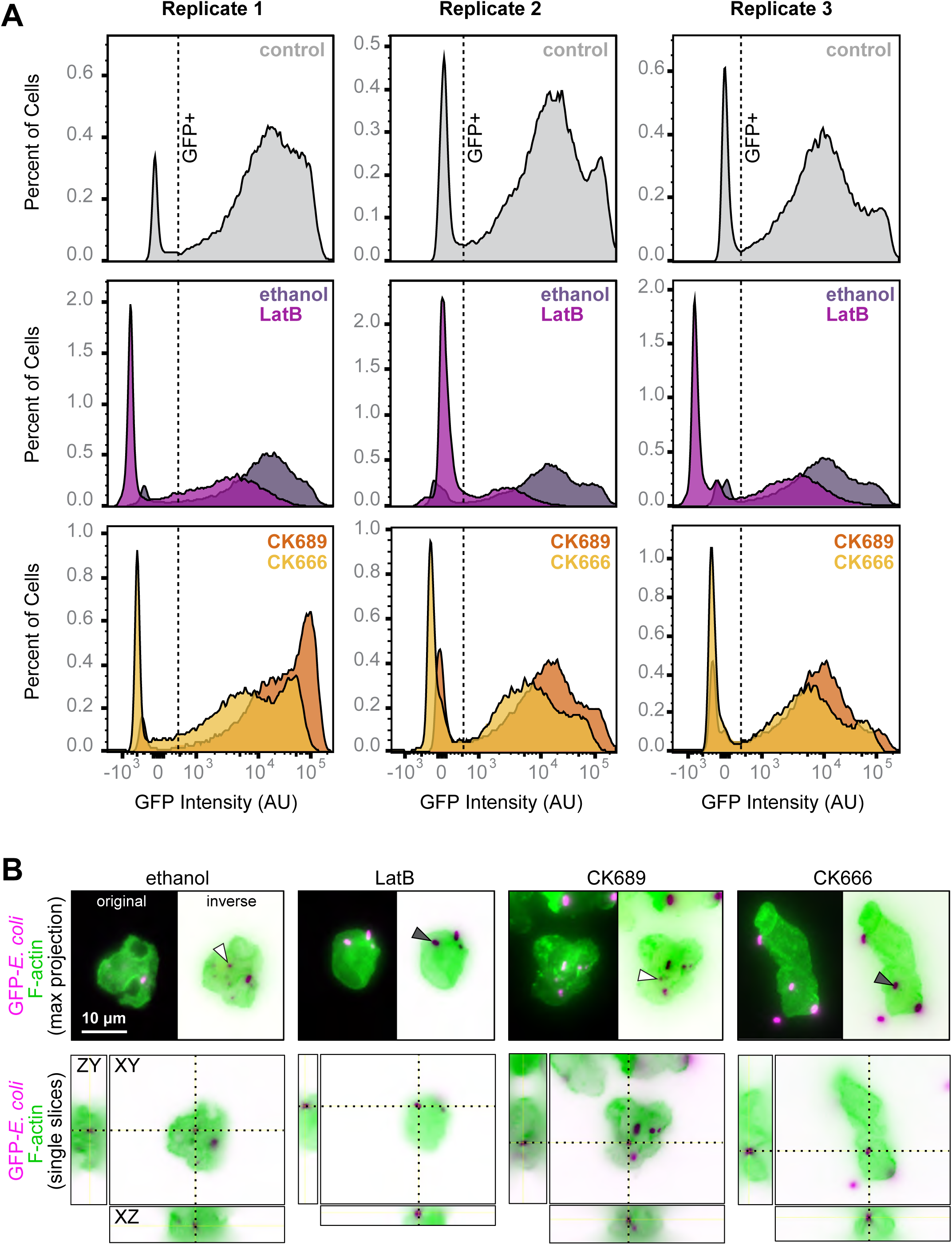
Latrunculin B and CK-666 each impair *Naegleria*’s phagocytosis. (**A**) NEGM cells were starved, treated with inhibitors or controls, and fixed as in Fig 6. Then, the GFP intensities of 30,000 cells per condition were measured by flow cytometry. Cells were initially gated to select only intact, single cells. Then, an untreated control from each replicate (top row) was used to create a gate for GFP+ cells (dashed line). Histograms display the percent of cells plotted against GFP intensities for ethanol- and LatB-treated cells (middle row), or CK-689- and CK-666-treated cells (bottom row). The histograms from replicate 2 are also shown in Figure 6, with y axes set to the same scale. (**B**) NEGM cells were starved and then mixed with GFP-expressing E. coli (shown in magenta) for 1 h. Cells were fixed and stained with Alexa-568 labeled Phalloidin to detect F-actin (green), and imaged using widefield fluorescence microscopy. Maximum intensity projections (top), and single xy, zy, and xz slices are shown intersecting non-intact (white arrowheads) and intact (gray arrowheads) bacteria associated with representative cells.

## Notes

### Competing Interest Statement

The authors have declared no competing interest.

